# Adaptive, maladaptive, neutral, or absent plasticity: Hidden caveats of reaction norms

**DOI:** 10.1101/2022.06.21.497038

**Authors:** Martin Eriksson, Alexandra Kinnby, Pierre De Wit, Marina Rafajlović

## Abstract

Adaptive phenotypic plasticity may improve the response of individuals when faced with new environmental conditions. Typically, empirical evidence for plasticity is based on phenotypic reaction norms obtained in reciprocal transplant experiments. In such experiments, individuals from their native environment are transplanted into a different environment, and a number of trait values, potentially implicated in individuals’ response to the new environment, are measured. However, the interpretations of reaction norms may differ depending on the nature of the assessed traits, which may not be known beforehand. For example, for traits that contribute to local adaptation, adaptive plasticity implies non-zero slopes of reaction norms. By contrast, for traits that are correlated to fitness, high tolerance to different environments (possibly due to adaptive plasticity in traits that contribute to adaptation) may, instead, result in flat reaction norms. Here we investigate reaction norms for adaptive versus fitness-correlated traits, and how they may affect the conclusions regarding the contribution of plasticity. To this end, we first simulate range expansion along an environmental gradient where plasticity evolves to different values locally and then perform reciprocal transplant experiments *in silico*. We show that reaction norms alone cannot inform us whether the assessed trait exhibits locally adaptive, maladaptive, neutral or no plasticity, without any additional knowledge of the traits assessed and species’ biology. We use the insights from the model to analyse and interpret empirical data from reciprocal transplant experiments involving the marine isopod *Idotea balthica* sampled from two geographical locations with different salinities, concluding that the low-salinity population likely has reduced adaptive plasticity relative to the high-salinity population. Overall, we conclude that, when interpreting results from reciprocal transplant experiments, it is necessary to consider whether traits assessed are locally adaptive with respect to the environmental variable accounted for in the experiments, or correlated to fitness.

## Introduction

Environmental conditions vary in time and space, and this can have variable consequences on species’ ranges and persistence (Bonier *et al.* 2007; Melbourne-Thomas *et al.* 2021; Rafajlović *et al.* 2022a, Rafajlović *et al.* 2022b). In general, the ability of organisms to survive and reproduce under a wide range of environmental conditions is considered to be of crucial importance for the long-term persistence of populations, not least under ongoing global climate change, which affects ecosystems on many levels (Parmesan 2006; Reusch 2014; Brown *et al.* 2016; Foden *et al.* 2019). One way for individuals to successfully cope with temporally and/or spatially changing environments is through locally adaptive phenotypic plasticity (e.g., Lande 2009; Chevin & Lande 2011; Reusch 2014; Levis & Pfennig 2016; Johansson et al. 2017; Bay et al. 2017; Enbody et al. 2021; Pazzaglia *et al.* 2021; Noer et al. 2022). Phenotypic plasticity is defined as the capacity of a single genotype to give rise to different phenotypes depending on the environment (Bradshaw 1965), and it can be locally maladaptive, neutral, or locally adaptive, resulting in negative, no, or positive consequences on individuals’ fitness, i.e., the ability to pass on genetic material to the next generation (Ghalambor 2007; Chevin *et al.* 2010; Velotta et al. 2018; Storz & Scott 2021).

The effect of the environment on an organism’s phenotype can be inferred experimentally through common garden and/or reciprocal transplant experiments. Specifically, in reciprocal transplant experiments, individuals that are native to two different environments are reciprocally transplanted to each other’s environment and a number of trait values are measured. Reciprocal transplants can be performed either during the adult or the juvenile life stage, depending on in which stage of development the plastic responses occur (Taborsky 2006; Ortega et al. 2017; Svensson et al. 2018; Delgado et al. 2020). The difference in phenotypes with respect to the environmental conditions is referred to as a reaction norm (Chevin *et al.* 2010). For individuals that express plasticity in an adaptive trait, we expect reaction norms in that trait to have significant non-zero slopes when the optimal phenotype in the two environments differ. Note that an adaptive trait is such that a change of value in this trait towards (or away from) the locally optimal phenotype increases (or decreases) individuals’ fitness. However, highly adaptive plasticity is expected to ‘optimise’ individuals' fitness in environments within the species' niche (Holt 2003), i.e., to keep the fitness approximately constant across environments. Therefore, if adaptive plasticity in traits that are under environmental selection is sufficiently high, it is expected that other traits that are strongly correlated to fitness (and yet selectively neutral with respect to the environment), would have essentially unchanged phenotypes in stressful environments (Reusch 2014). The most obvious example of such a trait would be fitness itself (de Vienne 2021). However, to accurately estimate fitness empirically is challenging (Reid et al. 2019, Alif et al. 2022). Fitness is particularly difficult to estimate in short-term experiments, which last shorter than the generation-time of an individual. In such cases, it is common to rely on indirect fitness proxies such as short-term survival, individual growth rates, or individual performance in traits that are assumed to be critical for survival (e.g., Wood et al. 2014, Johansson et al. 2017, Rugiu et al. 2021), although these could, in fact, be poor indicators of long-term fitness (Lailvaux et al. 2010; see also a discussion in Bonser 2021). In this study, we use the term *fitness-indicator traits* to refer to traits that are indicators of true fitness.

The adaptive maintenance of functional phenotypes in different environments is referred to as *phenotypic buffering* (Bradshaw 1965). As emphasised by Reusch (2014), phenotypic buffering complicates the interpretation of reaction norms with respect to the involvement of locally adaptive plasticity. For example, reaction norms for fitness-indicator traits undergoing phenotypic buffering are expected to be flat (i.e., absence of significantly non-zero slopes), according to the definition above, which suggests an absence of plasticity in the fitness-indicator traits. Therefore, one may incorrectly conclude that because plasticity in the measured trait is absent, *adaptive* plasticity is absent, although plasticity in some other (unobserved) adaptive trait may, in fact, be very high. While this problem has been verbally pointed out by Reusch (2014), it has not, to our knowledge, been formally theoretically studied.

To illustrate how reciprocal transplant experiments can lead to different conclusions depending on what trait is measured, we consider the isopod *Idotea balthica* (Pallas, 1772), which successfully colonized the brackish-water Baltic Sea (refer to Figure 1 in Björck (1995) for a map of the region) from the marine environment (Leidenberger *et al.* 2012) after the opening of the sea about 8000 years ago (Björck 1995). In this paper, we exclude the transition zone, *sensu* Snoeijs-Leijonmalm *et al.* (2017), consisting of the Kattegat and the Danish Straits, in our definition of the Baltic Sea, unless otherwise stated. Outside the Baltic Sea, under typical marine salinity conditions, *I. balthica* individuals are iso-osmotic at a salinity of 35 practical salinity units (psu) and maintain hyperosmotic body fluids in lower salinity with the use of Na^+^/K^+^-ATPase ion pumps located in the pleopods (Postel *et al.* 2000). However, individuals from the typical marine environment cannot maintain homeostasis when placed in 6 psu water where they rapidly die due to the osmotic stress (Hørlyck 1973). By contrast, *I. balthica* that are native to the brackish Baltic Sea conditions seem to do well in salinities down to 3 psu (Leidenberger *et al.* 2012). Furthermore, salinity does not have any significant effect on oxygen consumption, food consumption, growth, or short-term survival (i.e., for 12 weeks, which is much less than the generation time of the species) for adult individuals native to salinities ranging from 5-10 psu throughout the species’ present distribution within the Baltic Sea (Wood *et al.* 2014). However, the survival of juveniles at the current range margin (Rugiu *et al.* 2021) is significantly reduced when salinity is lowered below 3 psu. Furthermore, throughout the Baltic Sea, there seems to be a reduced survival when salinity is lowered simultaneously with an increased temperature (mimicking future climate conditions; Rugiu *et al.* 2018). Interestingly, and in contrast to the Baltic Sea populations, a population outside the Baltic Sea, just to the north of the Öresund strait, did not exhibit any significant decrease in survival under future climate conditions with increased temperature and decreased salinity (Rugiu *et al.* 2018).

**Figure 1:**
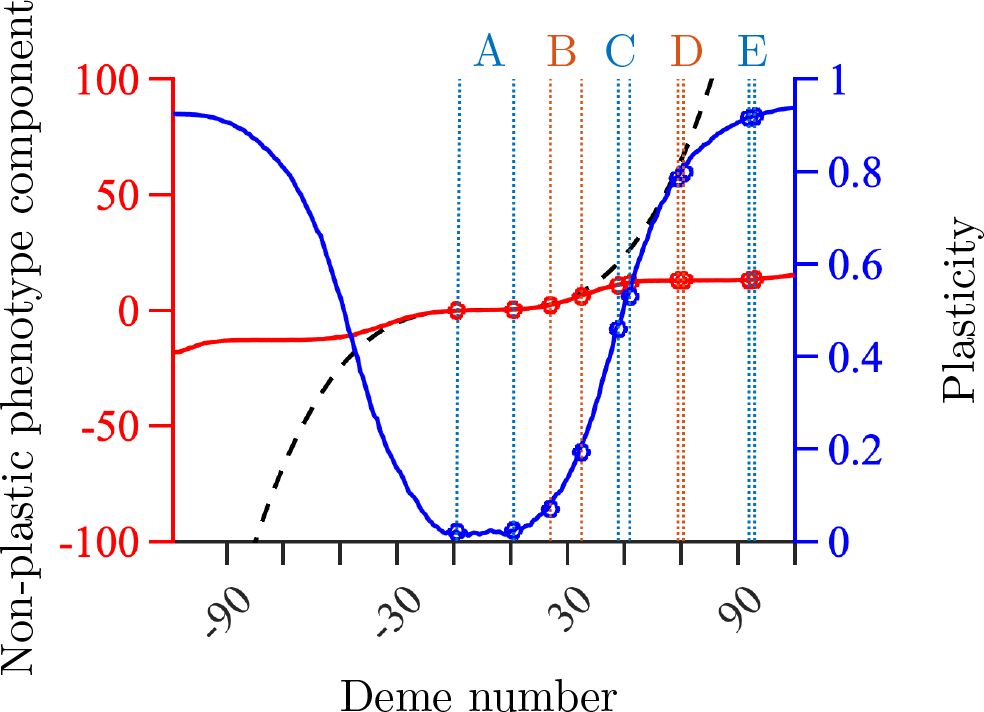
Spatial variation in the local average non-plastic component of the phenotype (red), the locally optimal adaptive phenotype (black dashed line) and local average plasticity (blue) at the end of the range expansion simulation in dependence of deme number. To better visualise the variability in the non-plastic component of the phenotype, the *y*-axis for the locally optimal adaptive phenotype and non-plastic component is truncated at ±100. The habitat consisted of M=220 demes, numbered from −109 to 110. The rings and the letters A, B, C, D, and E denote the experiments and the corresponding sampling locations for the experiments listed in Table 2 (with deme A1 to the left of A2 for experiment A, and similarly for the remaining experiments). Remaining parameter values: *K* = 100, *r_m_* = 2, *V_S_* = 2, *μ* = 10^−6^, *L* = 799, 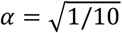, *β* = 2/*L* = 0.0025, *σ* = 1, *γ* = 0.5, and *δ* = 0.5 (Table 1).

The finding by Wood *et al.* (2014), that the expression of many traits in *I. balthica* are uniform throughout the Baltic Sea, seems to contrast with the results by Rugiu *et al.* (2018, 2021), who found that the populations inside the Baltic Sea are more sensitive to environmental changes than populations outside the Baltic Sea (north of the Öresund). Is this because there is a hard limit for plastic responses at approximately 3 psu, or is the population southeast of the Öresund (hereafter the *low-salinity population*) less tolerant to salinity changes than the population in the Kattegat and Skagerrak (hereafter the *high-salinity population*) and mainly adapted locally through the (non-plastic) genetic component of the phenotype, or is there an issue in how the reaction norms have been interpreted? The answer might lie in the nature of the measured traits, i.e., whether a trait is adaptive, or a fitness proxy (commonly assessed in short-term experiments). For example, an increase in metabolic rate could be adaptive if it would be correlated with higher food ingestion rates, growth, fecundity, and offspring survival. However, higher metabolic rates associated with less ingestion and growth would be a sign of stress and non-maintenance of function, in which case the trait could rather be considered fitness-indicative. In the former case, a non-zero reaction norm slope for metabolic rate would imply adaptive plasticity in this trait, whereas in the latter scenario, a zero-slope reaction norm would imply an adaptive response (with adaptive plasticity in some other unobserved trait under environmental selection) that acts to maintain fitness under novel conditions. Thus, it is often important to measure multiple traits simultaneously during reciprocal transplant experiments. However, even if this is done, the connection between trait values and fitness can remain unclear unless more information about fecundity and survival is available. Notably, additional clues can be obtained from theoretical models where information about fitness and the adaptive trait is readily available and can be used to produce expectations for reaction norms, taking into account different extents of local plasticity in the adaptive trait. As we show here, such theoretical models can help interpreting empirical data from reciprocal transplant experiments.

**Table 1:**
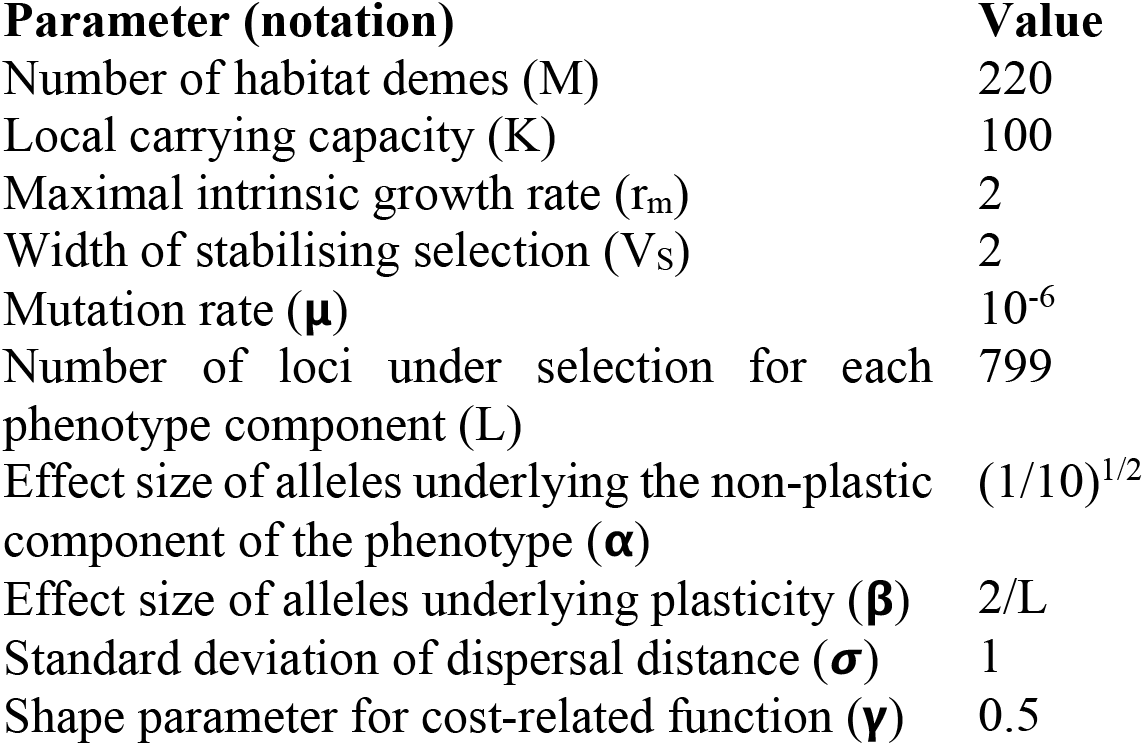
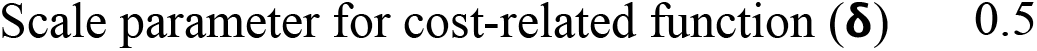
List of parameter values and notations used throughout.

| **Parameter (notation)** | **Value** |
| --- | --- |
| Number of habitat demes (M) | 220 |
| Local carrying capacity (K) | 100 |
| Maximal intrinsic growth rate ($r_m$) | 2 |
| Width of stabilising selection ($V_s$) | 2 |
| Mutation rate ($\mu$) | $10^{-6}$ |
| Number of loci under selection for each phenotype component (L) | 799 |
| Effect size of alleles underlying the non-plastic component of the phenotype ($\alpha$) | $(1/10)^{1/2}$ |
| Effect size of alleles underlying plasticity ($\beta$) | 2/L |
| Standard deviation of dispersal distance ($\sigma$) | 1 |
| Shape parameter for cost-related function ($\gamma$) | 0.5 |

**Table 2:**
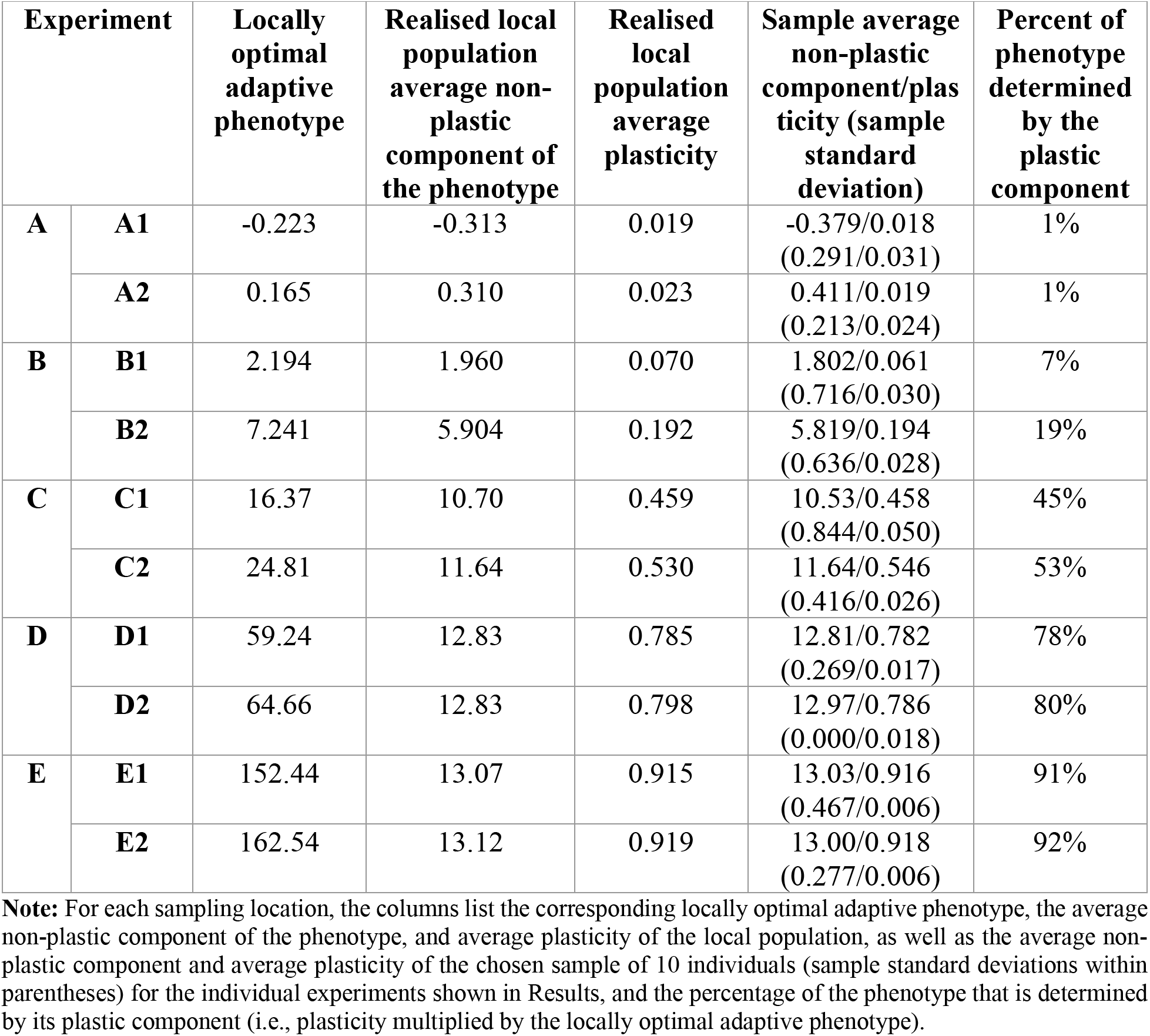
Sampling locations for the reciprocal transplant experiments.

| Experiment |  | Locally optimal adaptive phenotype | Realised local population average non-plastic component of the phenotype | Realised local population average plasticity | Sample average non-plastic component/plasticity (sample standard deviation) | Percent of phenotype determined by the plastic component |
| --- | --- | --- | --- | --- | --- | --- |
| A | A1 | -0.223 | -0.313 | 0.019 | -0.379/0.018<br>(0.291/0.031) | 1% |
|  | A2 | 0.165 | 0.310 | 0.023 | 0.411/0.019<br>(0.213/0.024) | 1% |
| B | B1 | 2.194 | 1.960 | 0.070 | 1.802/0.061<br>(0.716/0.030) | 7% |
|  | B2 | 7.241 | 5.904 | 0.192 | 5.819/0.194<br>(0.636/0.028) | 19% |
| C | C1 | 16.37 | 10.70 | 0.459 | 10.53/0.458<br>(0.844/0.050) | 45% |
|  | C2 | 24.81 | 11.64 | 0.530 | 11.64/0.546<br>(0.416/0.026) | 53% |
| D | D1 | 59.24 | 12.83 | 0.785 | 12.81/0.782<br>(0.269/0.017) | 78% |
|  | D2 | 64.66 | 12.83 | 0.798 | 12.97/0.786<br>(0.000/0.018) | 80% |
| E | E1 | 152.44 | 13.07 | 0.915 | 13.03/0.916<br>(0.467/0.006) | 91% |
|  | E2 | 162.54 | 13.12 | 0.919 | 13.00/0.918<br>(0.277/0.006) | 92% |

**Note:** For each sampling location, the columns list the corresponding locally optimal adaptive phenotype, the average non-plastic component of the phenotype, and average plasticity of the local population, as well as the average non-plastic component and average plasticity of the chosen sample of 10 individuals (sample standard deviations within parentheses) for the individual experiments shown in Results, and the percentage of the phenotype that is determined by its plastic component (i.e., plasticity multiplied by the locally optimal adaptive phenotype).

This paper has two aims. Our first aim is to understand when and how measurements of different traits result in different interpretations of reaction norms. To this end, we simulate reciprocal transplant experiments on simulated data. Importantly, we do not constrain our analysis to strictly ‘optimal’ phenotypic buffering *sensu* Reusch (2014). Instead, our simulated reciprocal transplant experiments account for population pairs sampled across a wide range of environmental conditions with low, intermediate or high local adaptive plasticity. We therefore find many cases where reaction norms in fitness-indicator traits have significant non-zero slopes (despite the involvement of locally adaptive plasticity in the adaptive trait). We refer to this as ‘sub-optimal phenotypic buffering’ because fitness is reduced, but not as much as it would have been in the absence of the plastic response. Based on the simulated reciprocal transplant experiments, we summarise a number of caveats associated with interpreting the norms of reaction.

Our second aim is to infer whether the low-salinity population of *I. balthica* has higher or lower tolerance to salinity changes than the high-salinity population, and whether the colonization of the Baltic Sea has occurred mainly through the evolution of non-plastic genetic adaptation to the low salinity environment or through the evolution of increased plasticity. To this end, we first performed reciprocal transplant experiments comparing the performance of *I. balthica* individuals from the north and the south of the Öresund strait (see Figure 1 in Björck (1995) where the Öresund is located to the east of Skåne in southern Sweden) in salinities of 16 and 8 psu, respectively. To our knowledge, reciprocal transplant experiments spanning the Öresund have not been performed to date in this species. Following this, we applied the conclusions from our model to interpret the reaction norms obtained from the empirical data in order to infer which of the two populations is likely to perform better in a wider range of salinities than the other.

## Methods

To illustrate the difference in reaction norms for the different kinds of traits and for populations with high and low plasticity, we performed reciprocal transplant experiments on simulated data, where the proportions of the plastic and non-plastic phenotypic components of the adaptive trait were known. To further illustrate how the interpretation of reaction norms can be altered depending on what trait is measured, we analysed empirical data from *I. balthica* sampled from the north and the south of the Öresund (i.e., from high- and low-salinity environments).

### Simulations of range expansion

Prior to simulating reciprocal transplant experiments, we performed simulations of a population’s range expansion over a habitat with spatially heterogeneous environmental conditions using the model presented in Eriksson & Rafajlović (2022). The model was not tailored to any specific species or habitat-specific environmental conditions. Instead, our main aim with the simulations was to create a spatially structured population suitable for illustrating how different extents of local plasticity and non-plastic local genetic adaptation can lead to different conclusions depending on what trait is measured. Thus, a conceptually general, rather than species-specific, model was used, parameterised to allow the population to evolve a wide range of plasticity values across the habitat. In our model, the population was assumed to expand its range across a habitat consisting of a one-dimensional array of demes. We modelled one quantitative trait under selection (i.e., the *adaptive trait*). The phenotypic value of the adaptive trait (hereafter the *phenotype* of the adaptive trait) was assumed to consist of a non-plastic and a plastic component that were allowed to evolve, and they approximately stabilised during the timespan simulated (100,000 generations). In the model, the local phenotypic optimum of the adaptive trait changed in space according to a spatially gradually steepening gradient (Figure 1, dashed black line). Note that one of the main factors that determine the locally optimal plasticity of the phenotype is the local steepness of the environmental gradient (Eriksson & Rafajlović 2022). Therefore, by choosing an environment with a spatially steepening gradient in the optimal phenotype, we allowed for a wide range of plasticity values to arise by evolutionary processes (rather than being set arbitrarily) within a single modelling framework. With an optimal phenotype that changes linearly in space, by contrast, a narrower range of plasticity values is typically obtained (Figure S1; Eriksson & Rafajlović 2022).

Generations were assumed discrete and non-overlapping. The population was assumed to be monoecious and mating was assumed to occur randomly (thus, self-fertilisation was possible, and it induced no fitness costs when it occurred). The individuals were diploid and the phenotype of the adaptive trait 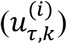 of individual *k* in deme *i* in generation *τ* was assumed to be determined by a plastic 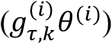 and a non-plastic 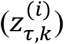 phenotype component (Eriksson & Rafajlović, 2022)

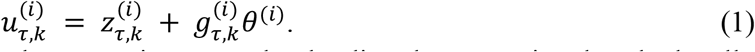

Here, the plastic component of the phenotype is assumed to be directly proportional to the locally optimal phenotype (*θ*^(*i*)^) and we refer to the coefficient of proportionality 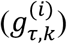 as *plasticity* (following Eriksson & Rafajlović 2022). Plasticity and the non-plastic component of the phenotype of the adaptive trait were polygenic, each determined by multiple bi-allelic loci with additive allele effects (with effect sizes ±*α*/2 for the non-plastic component of the phenotype and effect sizes ±*β*/2 for plasticity). The effect of the different parameters on the evolution of plasticity was extensively studied in Eriksson & Rafajlović (2022), and will not be considered in detail here. Instead, for the purpose of this study, we make use of the findings of Eriksson & Rafajlović (2022) to parameterise the model in such a way to 1) allow the population to occupy the entire habitat (e.g., by choosing allelic effect sizes and the number of loci appropriately) and 2) to produce a spatially heterogeneous pattern in plasticity with the average plasticity ranging from approximately 0 to approximately 1 in different parts of the habitat. The specific parameter values are listed in Table 1.

In the model, the fitness 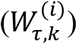 of individual *k* in deme *i* in generation *τ* was assumed to depend on the phenotype 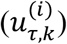 of the individual, the locally optimal phenotype (*θ*^(*i*)^), the local population size 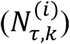, and (potentially costly) plasticity (Eriksson & Rafajlović, 2022):

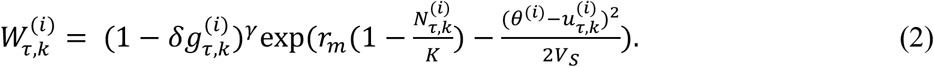

In equation (2), the term 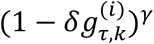 stands for the relative reduction in fitness due to the direct cost of plasticity, i.e., it does not include any fitness reduction due to potential maladaptive plasticity (hereafter this term is referred to as the cost-related function). The effect of maladaptive plasticity on fitness is accounted for by the last term of equation (2). The parameters *δ* and *γ* stand for the scale- and shape-parameter of the cost-related function, respectively (details in Eriksson & Rafajlović 2022). Although the empirical evidence for direct costs of plasticity is scarce (Murren et al. 2015, Snell-Rood & Ehlman, 2021), some costs or limits to plasticity must inevitably exist because, otherwise, plasticity would always outcompete non-plastic adaptation (Chevin & Lande 2011, Eriksson & Rafajlović 2022). We note that the results would be qualitatively similar if the cost-related function was set to 1 in the model, but unreliable environmental cues for the plastic response were used (whereas environmental cues were reliable in the present model). This is because unreliable environmental cues generate an implicit cost to plasticity by causing a mismatch between the phenotype generated by plasticity and the actual phenotypic optimum of the environment during selection (Tufto 2000). The parameters *r_m_*, *K*, and *V_S_* in equation (2) denote the maximal intrinsic growth rate, the carrying capacity, and the width of stabilising selection, respectively. All the parameters *δ*, *γ*, *r_m_*, *K*, and *V_S_* were assumed to be constant in time and the same for all individuals in all demes. The lifecycle was modelled as follows: Mating, recombination, mutation, formation of zygotes, death of adults, and dispersal of juveniles (details in Appendix A; see also Eriksson & Rafajlović (2022)). Recombination occurred freely between any pair of loci. Dispersal was assumed to occur according to a discretized Gaussian dispersal function (Eriksson & Rafajlović 2021) with standard deviation *σ* = 1. The realised gene flow between demes was an evolving property of the model, governed by dispersal, the environmental selection, mutation, and drift.

The parameter values examined here (Table 1) correspond to those used in Eriksson & Rafajlović (2022) except that *r_m_* = 2 here (the maximal intrinsic growth rate was larger than in Eriksson & Rafajlović (2022) to ensure families with tens of offspring in the *in-silico* experiments, which would be in line with the high reproductive rate of *I. balthica*). At the end of the simulations, a gradient in plasticity formed, with plasticity ranging from approximately 0 in the centre of the habitat, where the environmental gradient was shallow, to approximately 1 in the edges, where the environmental gradient was very steep, i.e., population average plasticity was strictly positive (Figure 1, blue line).

### Simulations of reciprocal transplant experiments

To simulate reciprocal transplant experiments, we sampled individuals from local populations obtained at the end of the range-expansion simulations described above. For each reciprocal transplant experiment, two different demes with different locally optimal adaptive phenotypes (and different contributions from the non-plastic and the plastic components to the phenotype; Figure 1) were chosen in such a way that qualitatively different results from the reciprocal transplant experiments were expected (Table 2). Due to the steep environmental gradient in the simulations, mainly relatively short-range transplants were performed (although longer-range transplants were also considered), and the pairs of demes were chosen in such a way to cover various combinations of plasticity differences in the different pairs of populations involved. Because the habitat was symmetric around the centre with respect to the horizontal axis, all samples, except sample A1, were taken from the right half of the habitat. From each deme, 10 individuals were sampled at random. Each of these individuals was assumed to have mated randomly within the local population before sampling. Since *r_m_* = 2, the number of offspring in the absence of density regulation (which is typically a good approximation for lab conditions) was large enough (i.e., on average 15 offspring for a well-adapted individual) allowing for large families in each experiment. Here we chose 10 offspring, uniformly at random, per sampled individual to reduce the effect of stochastic fluctuations with respect to individual family sizes. Thus, 10 families, each consisting of 10 half-siblings (with a possibility of some being full-siblings) were used in the simulated reciprocal transplant experiments.

The simulated reciprocal transplant experiments were performed by choosing uniformly at random 5 offspring per family to remain in their native environment, while the remaining offspring were transplanted into the new environment.

To account for stochasticity, we performed 1,000 realisations for each sampling location.

### Analysis of simulation results

From each reciprocal transplant experiment, we extracted reaction norms for the adaptive trait by measuring its realised phenotype in the native and the new environment in a given experiment. Furthermore, reaction norms were obtained for a trait that was assumed to have a trait value correlated to fitness (hereafter the *fitness-indicator trait*). The fitness-indicator trait was not explicitly modelled. Instead, its phenotype was assumed to be regulated by how locally well-adapted an individual is (e.g., through altered signalling pathways in new environments). Note that, as a consequence of this assumption, the fitness-indicator trait exhibits plasticity whenever fitness differs between the native and the new environment. To simplify the analysis, we defined the phenotype of the fitness-indicator trait to be equal to the phenotype-dependent component of fitness, obtained by setting 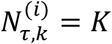 in equation (2), yielding

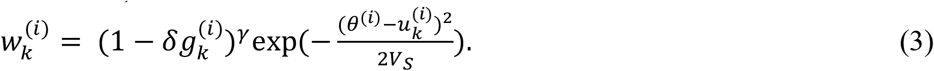

The notation in equation (3) is the same as in equation (2), except that here we omit the index for generations to emphasise that only one generation is considered during the reciprocal transplant experiment. Since fitness was assumed to be the main variation-causing factor in the fitness-indicator trait among individuals in the model, we used fitness itself as the fitness-indicator trait for simplicity. In this way, any plasticity in the fitness-indicator trait is strictly neutral in our model, which is a good approximation for a trait that is strongly correlated to fitness and either evolves neutrally or has an optimum value that is uniform among the different environments.

To test for differences in the mean phenotype between the native and the new environment for each pair of sampled populations, we used the Kruskal-Wallis test (Kruskal & Wallis 1952) followed by Bonferroni’s *post hoc* test (Miller 1981) for multiple comparisons. The Kruskal-Wallis test was preferred over an ANOVA to avoid the requirement of normally distributed residuals. Empirical data was analysed similarly (see below). Effect sizes for differences in the mean phenotypes were assessed using Glass’ Δ estimator (Glass et al. 1981).

To assess genetic differences between populations sampled from different demes, principal component analysis (PCA) was performed on the genetic similarity matrix, accounting for all simulated loci (underlying both the plastic and the non-plastic component of the phenotype).

The insights from the simulated reciprocal transplant experiments (described above) guided our interpretation of empirical data from salinity trial experiments of *I. balthica* (explained next).

### Empirical salinity trial experiment

During summer of 2018, *Idotea balthica* individuals from two different locations, spanning the steepest part of the salinity gradient of the Baltic Sea (including the transition zone), were used in a salinity trial experiment. The distance between the sampling locations was much larger than the typical dispersal distance (excluding potential rare rafting events), meaning that there was presumably little gene flow between the two populations (whereas this was not the case for all our simulated reciprocal transplant experiments; Figure 1, and see “Direction of significant reaction norms and their effect sizes” in Results). Adult isopods were collected in Vejbystrand (the high-salinity population; N 56.32°, E 12.76°; native mean salinity 16 psu) and in Svarte (the low-salinity population; N 55.43°, E 13.72°; native mean salinity 8 psu; salinity data are 0-6 m depth averages across 1995-2004 from the Rossby Centre Oceanographic circulation model (Meier, Döscher, & Faxén, 2003)) and were reciprocally transplanted into their native and the alternative (higher or lower) salinity (n≈75 per treatment for the high-salinity population, n≈50 per treatment for the low-salinity population). Isopods were kept in individual flow-through tanks for a period of 2 months, while fed their native diet, the brown alga *Fucus vesiculosus*. To avoid differences in palatability and metabolite composition of the food, which have been found to vary by geographic location (Nylund et al. 2012, Milec et al. 2022), all algae were collected near the Tjärnö Marine Laboratory (N 58.88°, E 11.14°), where the experimental work was conducted. In this way, we ensured that it was only the salinity that varied between the two environments.

At the onset of the experiment, the algae were separated into equally large pieces and placed in individual jars with or without isopods. We chose to focus on two commonly measured physiological traits for the examination of plasticity: Food consumption and respiration (as a proxy for metabolic rate). Grazing rates were monitored by weighing algae at the start and end of the experiment (carried out at 13 °C). The ungrazed algae pieces were used as controls for growth/water accumulation in tissues. After the 2-month period, oxygen consumption rates were measured for each individual isopod. Each individual was placed in a 5 mL glass vial and incubated at 6 °C in the dark for one hour. Before and after the incubation period, the oxygen concentration was measured every 15 seconds for a total of three minutes with a PreSens optical oxygen sensor. We present the mean values of these measurements. After the oxygen consumption measurements, the isopods were weighed and killed by decapitation and immediately placed in 95 % ethanol. Genetic analyses were used to investigate population genetic differences among the two collecting locations, as well as to ensure that the reciprocal transplant was random with regard to the genetic makeup of the isopods (see below).

### Analysis of empirical data

The grazing and respiration rate data were standardized by the natural logarithm of the weight of the isopods, as both response variables were correlated to weight. The two populations were then analysed for a significant effect of transplant salinity using the Kruskal-Wallis test followed by Bonferroni’s *post hoc* test for multiple comparisons. The effect sizes for differences in the mean phenotypes were assessed using Glass’ Δ estimator. In addition, we note that the grazing/weight values were normally distributed, whereas the respiration/weight values were normally distributed after transforming the raw data using the log10 transform. Thus, in addition to the Kruskal-Wallis test, we also performed a two-way ANOVA on the empirical data to test for differences between the two populations in terms of non-plastic differentiation, the common effect of the environment on the two populations, and gene-environment interactions. After examination of homoscedasticity and normal distribution of the residuals for the grazing/weight values and the log10-transformed respiration/weight values, three outliers were detected using Grubb’s test and removed before the two-way ANOVA was applied (note that outliers were only removed prior to the ANOVA and not the Kruskal-Wallis test).

### 2b-RAD sequencing

DNA was extracted from all isopods collected in the field for the purpose of the experiment using a Qiagen Blood & Tissue kit, after which 2b-RAD libraries were prepared using a modified version of the protocol designed by Mikhail Matz (https://github.com/DeWitP/BONUS_BAMBI_IDOTEA). 2b-RAD fragments were extracted from raw data by removing adapter sequences and filtering for the BcgI recognition site, after which data were quality filtered using a cut-off Q > 20. Reads were then mapped to the genome sequence using bowtie2 (Langmead & Salzberg 2012).

Individuals used in the empirical experiment were genotyped for 33,774 single nucleotide polymorphisms (SNPs), previously used by De Wit et al. (2020) using bcftools mpileup (https://samtools.github.io/bcftools/bcftools.html). After filtering to remove sites with >30 % missing data, 28,507 remaining polymorphic sites were used for principal component analysis, in order to examine the degree of genetic differentiation between the two populations.

## Results

### Reaction norms from simulated data

In the following, we present results obtained from simulated reciprocal transplant experiments for the sampling locations presented in Table 2.

As expected, with low plasticity and a small difference in locally optimal adaptive phenotype between the two sampling locations (A1 and A2), in a majority of realisations (96%) there was no significant phenotype difference in the adaptive trait between individuals in the native and the new environment, both for individuals that were native to deme A1 and for individuals that were native to deme A2. By contrast, in the majority of realisations (86%), there was a significant difference in the fitness-indicator trait both for individuals sampled from deme A1 and for individuals sampled from deme A2. These findings are further supported by small effect sizes (Δ) for the phenotype differences between the environments for the adaptive trait (Δ < 0.1) but large effect sizes for the fitness-indicator trait (Δ > 1). A typical simulation result for the reaction norms is shown in Figure 2A. In this simulation, there was a clear genetic differentiation between the population that was native to deme A1 and the population native to deme A2 (Figure S2A), mainly because the non-plastic component of genetic adaptation differed between the two populations.

**Figure 2:**
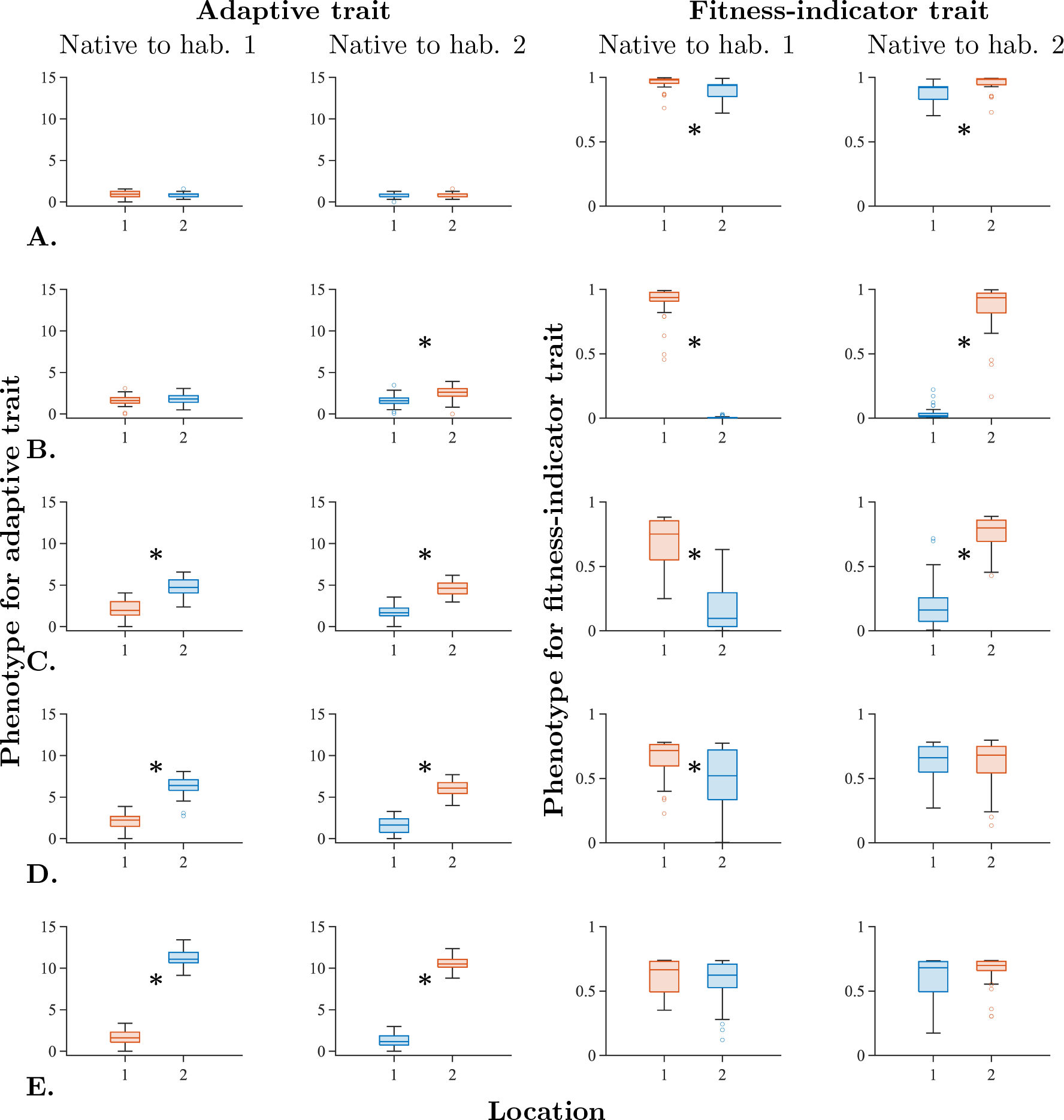
Boxplots of phenotypes from simulated reciprocal transplant experiments denoted by A-E in Table 2. Measured traits are the adaptive trait, z (column 1 and 2) and the phenotype-dependent component of fitness (column 3 and 4). Individuals that are native to environment Y1 are in column 1 and 3, individuals that are native to environment Y2 are in column 2 and 4. Here, Y=A-E in row 1-5, respectively. Kruskal-Wallis test *p*-values are: A: >0.99, 0.76, 4.1 · 10^−4^, 3.8 · 10^−10^; B: 0.74, 1.2 · 10^−7^, 5.0 · 10^−25^, 7.9 · 10^−11^; C: 8.9 · 10^−14^, 3.2 · 10^−18^, 6.4 · 10^−15^,1.1 · 10^−15^; D: 4.3 · 10^−16^, 5.8 · 10^−18^, 0.0011, >0.99; E: 8.2 · 10^−18^, 1.3 · 10^−16^, >0.99, 0.59. Significant (p < 0.05) differences are denoted with an asterisk in the plots. Glass’ *Δ* effect sizes are: A: −0.002, −0.067, −1.150, −1.333; B: 0.370, −1.092, −8.049, −5.074; C: 2.382, −3.698, −2.666, −4.489; D: 4.465, −4.849, −1.083, 0.067; E: 9.957, −13.735, −0.347, −0.564. The lines inside the boxes denote the median of the data and the boxes span between the upper and lower quartiles (75th and 25th percentiles). The whiskers indicate the maximum and minimum of the non-outlier data. Outliers (denoted by rings) were computed using the interquartile range (i.e., points above the upper quartile + 1.5 times the distance between upper and lower quartile or below the lower quartile - 1.5 times the distance between the upper and lower quartile were considered outliers). Trait values in the native environments are denoted by red box plots and trait values in the new environments are denoted by blue box plots. All phenotypic values for the adaptive trait are translated so that the minimum value measured in each experiment is zero.

The most common outcome of the reciprocal transplant experiments for sampling locations B1 and B2 was that there was no significant difference in phenotype of the adaptive trait between the native and the new environment for individuals that were native to deme B1, but a significant difference, with large effect size, between native and new environment for individuals that were native to deme B2 (Figures 2B, S1B). This outcome occurred in 55% of the realisations, and in the other 45% of the realisations, none of the two populations had a significant difference in the adaptive trait. For the fitness-indicator trait, there was in all 1000 realisations a significant difference with large effect size between the phenotype in the new and the native environment, both for individuals that were native to deme B1 and for individuals that were native to deme B2. However, the reduction in fitness was less severe for individuals that were native to deme B2 than to B1. As for sampling locations A1 and A2, there was a clear genetic differentiation between the two populations (Figure S2B).

For intermediate plasticity (sampling locations C1 and C2), there were significant phenotype differences with large effect sizes for both traits and both populations in all 1000 realisations, and the populations native to the two different environments were genetically differentiated (Figures 2C, S1C). However, the reduction in fitness was smaller than for demes B1 and B2, despite a larger difference in locally optimal adaptive phenotype.

When there was almost no difference in the non-plastic phenotype component of the adaptive trait, but the plasticity was high and differed between the two populations (sampling locations D1 and D2), we observed a significant difference with large effect size in the adaptive trait between the native and new environment for both populations in all 1000 realisations. For the fitness-indicator trait, we observed a significant reduction in fitness with typically large effect size in the new environment for individuals that were native to deme D1 but not for individuals that were native to deme D2 in 40% of the realisations (Figures 2D, S1D). In 32% of realisations, we found that none of the populations had significant differences in phenotypes for the fitness-indicator trait. The remaining two possibilities, namely that both differences were significant, or that only the population native to D2 but not the one native to D1 had significantly non-zero reaction norms, each occurred in 14% of the realisations. In the simulation result shown in Figures 2D, S1D (a single randomly chosen realisation) there was a less clear separation between the two populations by PCA than for populations sampled at locations A1 and A2, B1 and B2, or C1 and C2, despite the relatively large difference in the locally optimal adaptive phenotype (Figure S2D; compare to panels A-C). Note that we did not have neutral loci in the model, which could contribute to differentiation between natural populations.

When plasticity was very high (>0.9) in both environments (sampling locations E1 and E2) and the difference in locally optimal adaptive phenotype between the two environments was large (Δ*θ* ≈ 10), phenotypes of the adaptive trait exhibited significant differences with very large effect sizes between the native and the new environment for both populations in all 1000 realisations. However, despite the large difference in the locally optimal adaptive phenotype between these demes, no significant differences in the fitness-indicator trait values occurred in 51% of the realisations (Figures 2E, S1E). In 22% of the realisations, a significant difference in the fitness-indicator trait values occurred only for the population native to deme D2, whereas in 18% of the realisations a significant difference in the fitness-indicator trait values occurred only for the population native to deme D1. None of the populations had significant differences in the fitness-indicator trait in 9% of the realisations. Typically, the effect sizes for the fitness-indicator trait were small to moderate in magnitude. For the simulation result shown in Figure 2E, there was no clear genetic differentiation between the two populations (Figure S2E). Typical results from the simulated reciprocal transplant experiments are summarised in Table 3.

**Table 3:** Summary of results from the simulated reciprocal transplant experiments.

|  | Significant phenotype difference in adaptive trait |  | Significant phenotype difference in fitness-indicator trait |  |
| --- | --- | --- | --- | --- |
|  | *Native to env. 1* | *Native to env. 2* | *Native to env. 1* | *Native to env. 2* |
| Experiment A:<br>Very low plasticity | No | No | Yes | Yes |
| Experiment B:<br>Low plasticity | No | Yes | Yes | Yes |
| **Experiment C:**<br>**Intermediate**<br>**plasticity** | Yes | Yes | Yes | Yes |
| **Experiment D:**<br>**High plasticity** | Yes | Yes | Yes | No |
| **Experiment E:**<br>**Very high**<br>**plasticity** | Yes | Yes | No | No |

Note: In all cases, plasticity is higher for individuals that are native to environment 2 than to environment 1. The table shows typical outcomes for the listed experiments (Experiment A-E). Due to randomness, the presented significances may not occur in every realisation.

For reciprocal transplant experiments across larger distances in our model (recall that the environmental gradient was steepening in space), the reaction norms for the fitness-indicator trait were always significant with large effect sizes (typically with *p* ≪ 10^−10^ and |Δ| ≫ 1; Figure S3). For the adaptive trait, the reaction norms were also typically significant, except for when plasticity in the native environment was very low (Figure S3 A). Note that fitness, in all cases, was extremely low in the new environment, which implies that individuals would not be able to reproduce there, although they may survive in short-term experiments (which could lead to an overestimation of the actual fitness in the experiments). In all cases, there was a clear genetic differentiation between the two populations (Figure S4).

Figure 2 indicates that the relative directions of significant reaction norms obtained in pairs of reciprocal transplants may differ between adaptive and fitness-indicator traits. For example, the adaptive trait value was higher in deme Y2 than in deme Y1 when Y=C, D or E in the realisations shown (but different samples from the same localities can give different results, as explained above). By contrast, the fitness-indicator trait value was higher in the native environment (regardless of whether the native environment is deme Y1 or Y2) than in the new environment when Y=A, B or C. We analyse this further next.

### Direction of significant reaction norms and their effect sizes

To further understand how the difference in local phenotypic optima and plasticity influence the reaction norms, including their significance, direction, and effect sizes, we simulated additional reciprocal transplant experiments with one (and the same) sampling location being involved in each experiment as the “focal” deme, whereas the second “alternative” deme varied among the experiments. The focal deme was either deme 0 (with an average plasticity of 0.02), or deme 20 (with an average plasticity of 0.06). The alternative deme was every second deme between deme 2 and deme 110, or between deme 22 to deme 110, respectively (see Figure 1 for deme numbers and an illustration of the habitat). Note that the population native to any alternative deme had higher plasticity on average than the population native to either focal deme (blue line in Figure 1). Reaction norms for the adaptive trait were typically in opposite directions (that is, the phenotype difference between foreign and native environments were of opposite sign), when individuals were transplanted from their respective native environments to the new environment, as expected (Figures 3A). The effect sizes of the phenotypic differences were large when the differences in locally optimal phenotype (Δ*θ*) between the focal and alternative environments were large enough (Figure 3A). However, the reaction norms for the adaptive trait had significant non-zero slopes only when both the local plasticity in the native environment and Δ*θ* were sufficiently large. Consequently, when the focal environment was deme 0, and Δ*θ* ≥ 5, the slopes of reaction norms for the adaptive trait were not significantly non-zero for individuals that were native to the focal environment (Figure 3C blue rings), whereas the reaction norms for the adaptive trait were significantly non-zero for individuals that were native to the alternative environment (Figure 3C red crosses). Note that, even when the slopes of the reaction norms were not significantly non-zero, their effect sizes could be large when Δ*θ* was large enough (Figure 3A). Conversely, when the focal environment was deme 20 (with higher plasticity than deme 0), the directions of the reaction norms for the adaptive trait were qualitatively the same as when the focal environment was deme 0 (Figure 3E), but the slopes of the reaction norms for individuals that were native to the focal environment were significantly non-zero when Δ*θ* ≥ 10 (Figure 3G blue rings) whereas the reaction norms were significantly non-zero when Δ*θ* ≥ 5 for individuals native to the alternative environment (Figure 3G red crosses).

**Figure 3:**
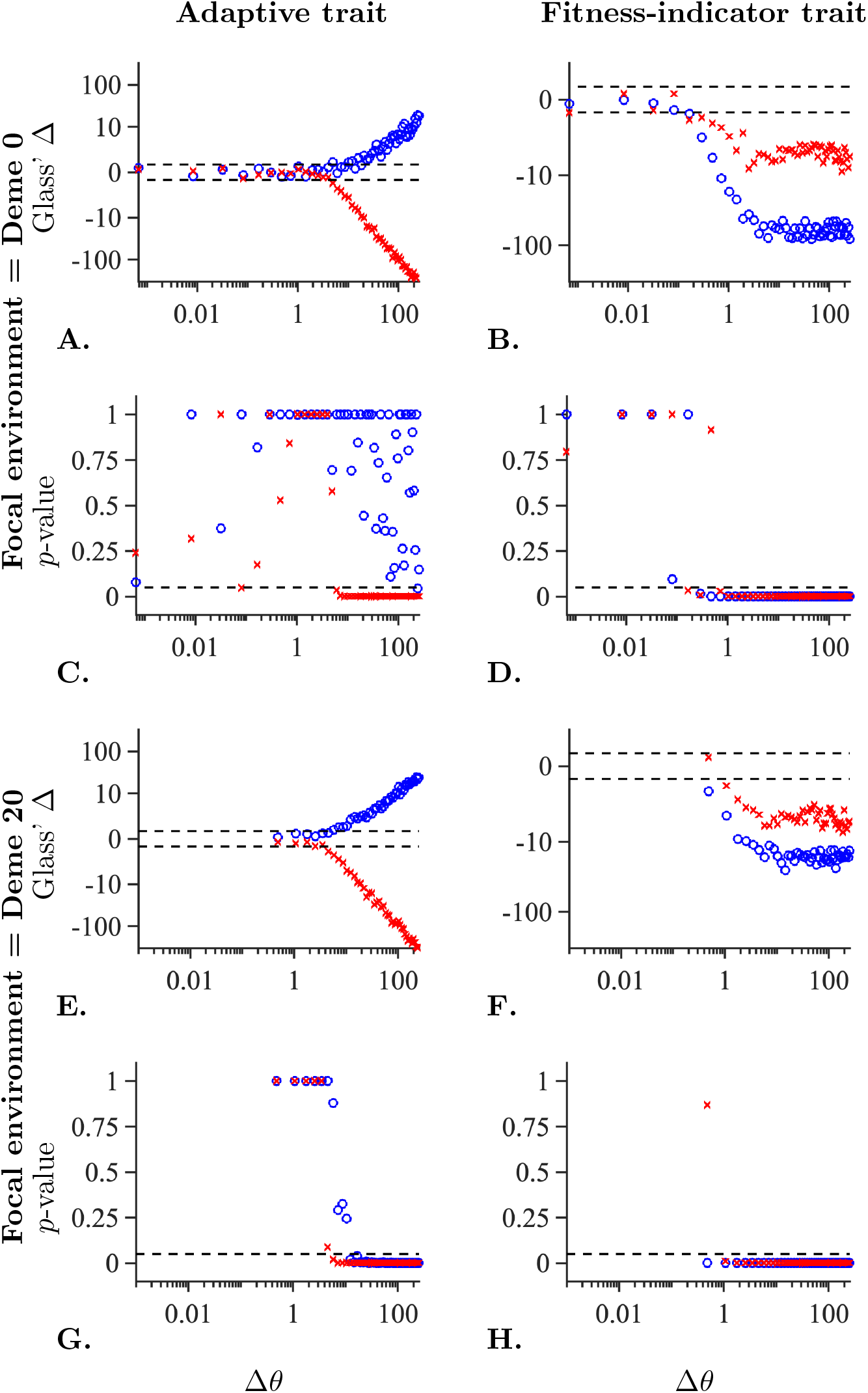
Plots of effect sizes with direction (A, B, E, and F) and *p*-values for non-zero slopes of reaction norms (C, D, G, and H) as a function of the difference in locally optimal adaptive phenotype (Δ*θ*) between the focal and the alternative environment for the adaptive trait (left column), or for the fitness-indicator trait (right column). Results are shown for focal deme numbered 0 (A-D) or 20 (E-H). The blue rings stand for reaction norms for individuals that are native to the focal deme (with low plasticity) whereas red crosses stand for reaction norms for individuals that are native to the alternative deme (where plasticity is higher than in the focal deme). The dashed lines indicate the line beyond which absolute values of effect sizes can be considered *large* (C, D, G, H) according to Cohen’s rules-of-thumb (Sawilowsky, 2009), or where *p*=0.05 for the Kruskal-Wallis test (C, D, G, H). Note a logarithmic scale on the *y*-axis in panels A, B, E, and F, and on the *x*-axis in all panels.

By contrast, the slopes of reaction norms for the fitness-indicator trait were typically in the same direction when individuals were moved from their native environment to a new environment (Figures 3B, 3F). The realised reaction norms were flat when the difference in the locally optimal adaptive phenotype between the focal and the alternative environment was too small in our simulations, that is, less than Δ*θ* = 0.1 when the focal environment was deme 0 (Figure 3D) and less than Δ*θ* = 1 when the focal environment was deme 20 (Figure 3H).

When Δ*θ* was kept constant and *both* populations were moved along the environmental gradient such that the plasticity in the native environment was varied, reaction norms for the adaptive trait were flat when plasticity was very low (Figure S5 A) whereas reaction norms for the fitness-indicator trait were flat when plasticity was high (Figure S5 B). The effect sizes for differences in the adaptive trait were large when plasticity was high (Figure S5 C). By contrast, the effect sizes for the differences in the fitness-indicator trait were very large when plasticity was low, but small when plasticity was high (Figure S5 D).

We additionally examined the norms of reaction in a model with a phenotypic optimum that changed linearly in space (rather than according to a steepening gradient). In the experiments with a fixed focal environment and a varying alternative environment in this model, the results were qualitatively similar to those in Figure 3. However, for the linearly changing optimum, the spatial variability of plasticity was smaller than in the model with a steepening environmental gradient (Figure S1). Consequently, the transplant results between the focal and alternative environments were more rarely asymmetric (i.e., with a significant non-zero slope of the norm of reaction for only one population) than in the model with a steepening gradient (Figure S6), although some asymmetry existed when the environmental gradient was shallower, owing to a stronger gradient in realised plasticity.

### Empirical data

For the population collected in the high-salinity environment (native salinity 16 psu), neither grazing nor respiration exhibited a statistically significant difference between the two experimental salinity treatments (Figure 4 A-B). For the population collected in the low-salinity environment (native salinity 8 psu), however, grazing rates were significantly lower (p=0.008; effect size Δ = −0.538) and respiration rates were higher (p=0.003; Δ = 0.521) in the high-salinity environment than in the native salinity (Figure 4 C-D). Two-way ANOVAs did not show significant differences in either grazing or respiration rate between the two populations (Tables S1-S2). However, there was a significant difference between the environments for grazing (Table S1), but not for respiration rate (Table S2). Furthermore, there were significant gene-environment interaction (GxE) effects in respiration rate and grazing. Bonferroni’s *post hoc* test revealed that the significant differences were between the native and the new salinity for individuals from the low-salinity population for both grazing and respiration rate. For respiration rate, there was also a significant difference between low-salinity and high-salinity individuals in a salinity of 8 psu (Table S2). The genetic data showed that the two populations are genetically differentiated (Figure 5).

**Figure 4:**
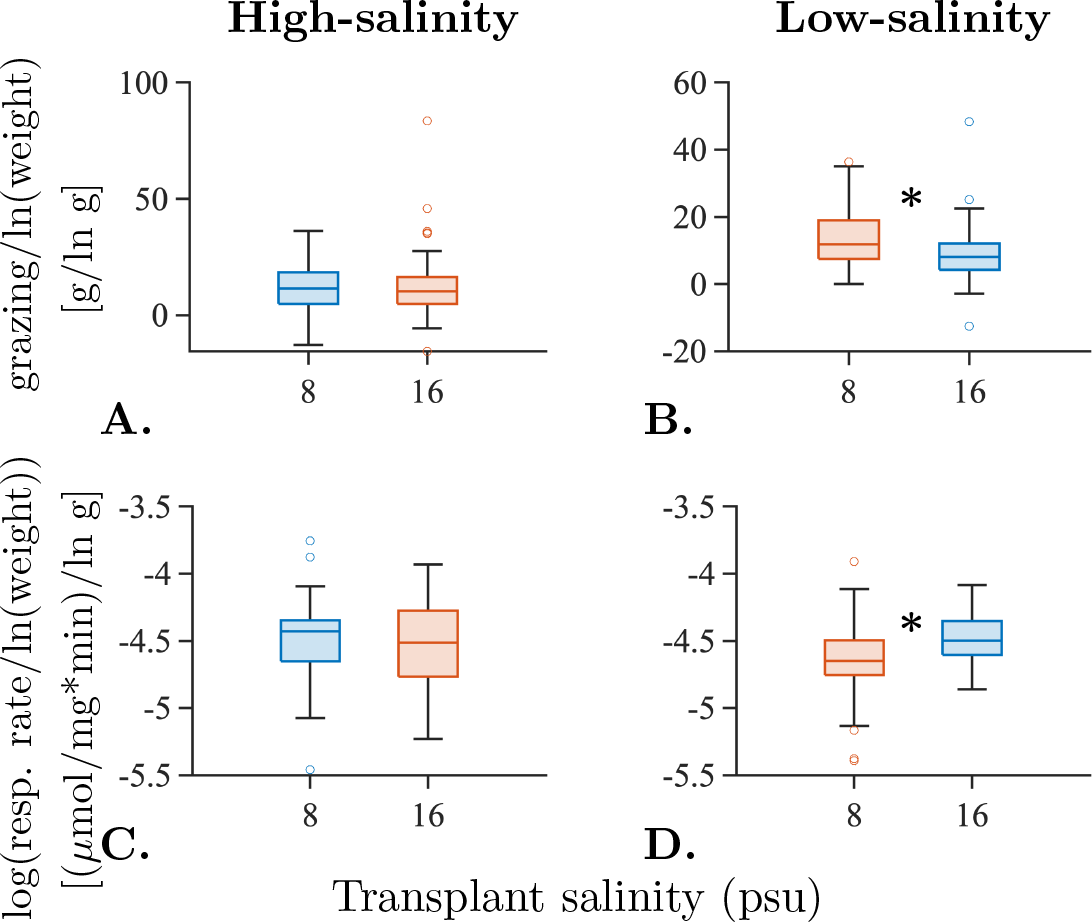
Boxplots of empirically measured phenotypic traits in reciprocally transplanted isopod *Idotea balthica* samples from the high-salinity environment (native salinity 16 psu; A and C) and the low-salinity environment (native salinity 8 psu; B and D). Measured traits are grazing per unit weight of the isopod (A and B), and the log10-transform of the respiration rate per unit weight of the isopod (C and D). Glass’ Δ estimates for effect sizes for transplants from the native to the new environment are: A: −0.054; B: −0.538; C: +0.123; D: +0.521. Kruskal-Wallis p-values are: A: >0.99; B: 0.049; C: >0.99; D: 0.047 (the two-way ANOVA p-values are 0.998, 0.0156, 0.267, and 0.0267, respectively). Significant (p < 0.05) differences are denoted with an asterisk in the plot. Trait values in the native environments are denoted by red box plots and trait values in the new environments are denoted by blue box plots.

**Figure 5:**
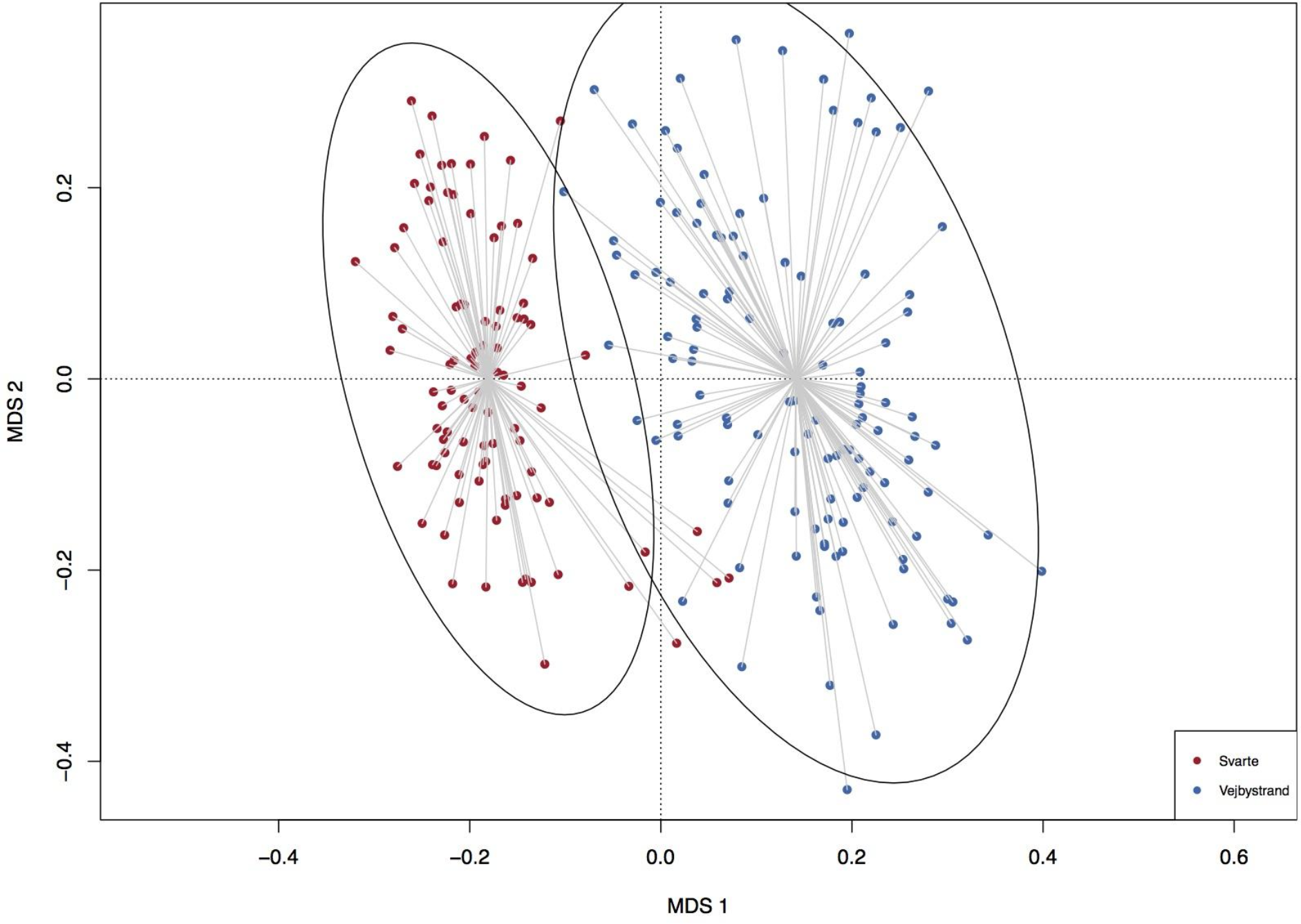
Principal component analysis of identity-by-state (IBS) distances, using 28,507 2b-RAD derived genetic loci. Low-salinity individuals are coloured in red, high-salinity in blue. Circles represent the 95% confidence intervals of the two experimental groups. Collecting location is a strong determinant of the genetic variation (PERMANOVA p < 0.001).

## Discussion

Reciprocal transplant experiments are often used to test how a population is affected by the environment. Non-zero reaction norms can be taken as either evidence for plasticity (e.g., Côte et al. 2019; Walter et al. 2021) or for local adaptation (e.g., Barton et al. 2020; Martin et al. 2021). However, the interpretation of reaction norms obtained in reciprocal transplant experiments may lead to opposite conclusions depending on whether the assessed trait is an adaptive trait or a fitness-indicator trait. To understand how these different kinds of traits may affect the interpretation of reaction norms, we performed reciprocal transplant experiments on simulated data. In the simulations, a population was allowed to expand its range across a finite habitat with a spatially varying optimal adaptive phenotype. The simulations were run for a sufficiently long period to allow all local populations to stabilise, whereupon reciprocal transplant experiments were performed and reaction norms were assessed. In addition, we performed reciprocal transplant experiments on two populations of *Idotea balthica* from the south and the north of the Öresund strait (see Figure 1 in Björck et al. (1995) for a map).

In the following, we summarise our theoretical findings regarding reaction norms from simulated reciprocal transplant experiments, assuming that both sampled populations have stabilised with respect to migration, selection, mutation, and drift (meaning that the average phenotypes of the populations in their home environments were close to the local optima). These findings are subsequently applied to the empirical data obtained from the reciprocal transplant experiments performed on the two populations of *I. balthica*. Although we do not know how close the two populations of *I. balthica* are to their optima, the arguments here work well as long as each population is better adapted to its native environment than to the environment to which it is transplanted.

### Reciprocal transplant experiments on simulated data

No significant differences between the trait values for the adaptive trait in the native and the new environment usually occurred for populations that had both negligible plasticity in the adaptive trait and a small difference between the phenotypic optima (although significant differences could sometimes occur by chance). By contrast, we found statistically significant non-zero slopes in reaction norms for the adaptive trait when plasticity in the adaptive trait was high and/or the difference in phenotypic optima was large.

In contrast to the adaptive trait, the slopes of reaction norms were significantly non-zero for the fitness-indicator trait when locally adaptive plasticity in the adaptive trait was too low to keep the difference in fitness between the new environment and the native environment statistically insignificant. That is, the slopes of reaction norms were significantly non-zero when phenotypic buffering of the fitness-indicator trait was sub-optimal. By contrast, when locally adaptive plasticity in the adaptive trait was high enough to keep fitness unchanged in the new environment relative to the native environment, i.e., when phenotypic buffering was optimal, there were no significant differences in the fitness-indicator trait between the two environments.

Note that, unless one of the populations is better adapted to the other population’s environment than to its own native environment (for example, if one population has very recently colonised its current habitat) the fitness in the non-native environment is expected to be either lower or equal to the fitness in the native environment. Thus, significant non-zero slopes of reaction norms for fitness-indicator traits have the same direction (i.e., the same sign of the phenotype difference between the new and the native environment) for both populations, unless one of the populations is better adapted to the native environment of the other population than to its own native environment. This may allow us to distinguish between fitness-indicator traits with suboptimal phenotypic buffering and adaptive traits with plasticity. Namely, when for both populations the reaction norms for a fitness-indicator trait are significantly non-zero and have the same direction (indicating a decrease in fitness in the non-native environment), potential plasticity in the adaptive trait is not sufficiently adaptive to maintain the locally optimal adaptive phenotype in the new environment.

When the reciprocal transplant experiments were carried out across a long enough distance (implying a large difference in phenotypic optimum in our model) we always observed a significantly lower fitness in the new environment compared to the native environment (because the phenotype was never determined solely by plasticity in our model; some non-plastic genetic adaptation was always involved). Thus, whether phenotypic buffering is optimal depends on the difference between the two environments: A population may experience optimal phenotypic buffering when the difference in optimal phenotype between the native and the new environment is small enough, but sub-optimal phenotypic buffering otherwise. Indeed, when local plasticity is higher in one of the environments than in the other, phenotypic buffering may be optimal for one population but sub-optimal for the other.

In our model, plasticity increases from the centre towards the edges of the habitat. Notably, however, environmental conditions in natural populations consist of both abiotic and biotic factors, e.g., species competition (Case & Taper 2000, Turcotte & Levine 2016), and the combined effect of these could lead to harsher conditions and/or steeper gradients in the central parts of the habitat than in the habitat edges (for example, if competition is stronger in central parts of the habitat), which could potentially promote the evolution of higher plasticity in the centre than in the edges. Elucidating the explicit role of biotic versus abiotic factors, and their interplay for the evolution of plasticity and non-plastic genetic adaptation is an interesting avenue for future work, both theoretically and empirically.

Recall that we here assumed that the fitness-indicator trait was equal to the phenotype-dependent component of fitness (which depends also on the cost of plasticity). However, assuming orthogonality between traits and free recombination between the loci underlying the traits, we expect that our conclusions hold also for any trait *f* that is a linear function of the phenotype-dependent component of fitness *w*, i.e., *f* = *a* + *bw*. Here, *a* and *b* are environment-independent trait components (or trait components having a spatially uniform optimal phenotype) with a polygenic basis, allowed to evolve over time, and *w* is the phenotype-dependent component of fitness, Eq. (4). Note that the coefficient *b* can take any real value, but for *b<0*, the trait *f* would be negatively correlated to *w*, whereas for *b=0* the trait would be uncorrelated to *w*. This is because the population average phenotype of a neutral trait orthogonal to an adaptive trait eventually evolves to a spatially constant value (Tomasini et al. 2022). The same is true for a trait under global selection with a spatially uniform optimal phenotype, as long as the global optimum is temporally constant (see, for example, Bisschop et al. 2020 for the effect of global selection on local adaptation *albeit* in a two-deme model). Further studies are needed to understand how potential correlations between traits that are fitness-indicators and traits that are adaptive affect the reaction norms of fitness-indicator traits.

### Interpretation of empirical data

We observed a statistically significant increase in respiration, measured as a proxy for metabolic rate, and a decrease in grazing activity in Baltic Sea *I. balthica* when transplanted from a native salinity of 8 psu to a salinity of 16 psu. By contrast, no significant differences in grazing activity or metabolic rate were apparent when *I. balthica* native to a salinity of 16 psu were transplanted to a salinity of 8 psu. Because flat reaction norms may suggest either very high locally adaptive plasticity (i.e., phenotypic buffering) or very low (or absent) locally adaptive plasticity, the possible conclusions with respect to the combinations of the types of traits measured (fitness-indicator or adaptive) are the following:

If both traits are adaptive, the results would suggest that the high-salinity population (native salinity 16 psu) does not have any significant plasticity with respect to salinity, whereas the low-salinity population (native salinity 8 psu) expresses plasticity in salinity tolerance. In this case, the plastic responses are expressed as a change in respiration rate and grazing activity. By contrast, if at least one of the traits is a fitness-indicator trait, the high-salinity population would have locally adaptive plasticity in some unmeasured adaptive trait, which keeps fitness unchanged when transplanted to the low-salinity environment (thus the fitness-indicator trait is phenotypically buffered *sensu* Reusch 2014). In this case, plasticity in the adaptive trait with respect to salinity is lower for the low-salinity population than for the high-salinity population (or the low-salinity population lacks plasticity in the adaptive trait with respect to salinity), and the low-salinity population, therefore, has a reduced fitness in the high-salinity environment. Note that it is sufficient that one of the two measured traits is a fitness-indicator to conclude that the low-salinity population of *I. balthica* has significantly lower fitness when transplanted to the high-salinity environment (and the conclusion remains the same regardless of which trait is a fitness-indicator). To determine which of the two possibilities is more likely (i.e., if both traits are adaptive, or at least one trait is a fitness-indicator trait), we consider what these traits imply for the organism, and what previous studies have found.

To our knowledge, there are no studies on the osmotic regulation in Baltic Sea populations of *I. balthica* (only of populations from Helgoland in the North Sea; Postel et al. 2000). However, the closely related species *Idotea chelipes* in the Baltic Sea (native salinity 7 psu) has been shown to osmoconform, and thus reduce metabolic rates (presumably due to less required ion pumping) at salinities above 11 psu (Łapucki & Normant 2008). Similarly, Normant & Lamprecht (2006) found that Baltic Sea individuals of the amphipod *Gammarus oceanicus* reduce their metabolic rates as well as feeding rates in higher salinities. These findings contrast with our result. The observed increase in metabolic rate for *I. balthica* should be accompanied with an increase in food intake as more energy is required, if the increased metabolic rate is locally adaptive. However, instead we observed a reduction in the amount of grazing. We further note that the grazing activity is the same in the native environments for both populations (Table S3), which further suggests that this may be a fitness-indicator trait that is phenotypically buffered with respect to salinity for the high-salinity population, but sub-optimally phenotypically buffered with respect to salinity for the low-salinity population. Combined, these observations suggest that the phenotypic changes observed when transplanting low-salinity population isopods to 16 psu seawater are non-adaptive with respect to salinity changes and that suboptimal phenotypic buffering occurs for the low-salinity population, at least for the grazing rate. By contrast, optimal phenotypic buffering with respect to reduced salinity seems to occur for the high-salinity population, suggesting that this population has higher locally adaptive plasticity with respect to reduced salinity than the low-salinity population has with respect to increased salinity. Importantly, however, it remains to be understood whether optimal phenotypic buffering for the high-salinity population would be retained if we included in the experiment also other environmental factors that differ between the sampling locations (and that may have been involved in adaptation). This is an important open question for future studies.

Interestingly, our data show that the respiration rate is the same in all environments except for the low-salinity population in its native salinity. By interpreting the respiration rate in the native environment as the locally optimal respiration rate, this finding would imply that respiration rate is an adaptive trait and that the low-salinity population, but not the high-salinity population, has locally adaptive plasticity in respiration rate. However, as the expression of many traits influences the metabolic rate and therefore also the respiration rate, the respiration rate is clearly correlated to multiple other traits, that may or may not be involved in adaptation to salinity. Genetic differentiation in these traits may cause the optimal respiration rate to differ between individuals from the low-salinity population and individuals from the high-salinity population, even when they are exposed to the same salinity level. Additionally, even if the plastic response in respiration rate is indeed adaptive, it could be counteracted by locally maladaptive plasticity in other traits that are also affected by salinity.

Note that the low-salinity population must still have a relatively high locally adaptive plasticity in the adaptive traits if metabolic rate and/or grazing are fitness-indicator traits, because metabolic rate and grazing are constant with respect to the salinity levels within the Baltic Sea (Wood *et al.* 2014). Thus, it seems likely that the low-salinity population either has a reduced salinity-related locally adaptive plasticity relative to the high-salinity population, but still high enough locally adaptive plasticity to maintain homeostasis within the entire Baltic Sea, and/or that it has an effective response across a different salinity range (e.g., between 3-15 psu, whereas the high-salinity population may tolerate a salinity range between 10-35 psu).

Our empirical results also suggest that there is some genetic differentiation between the high- and low-salinity populations. A recent study (De Wit *et al.* 2020) also highlighted a genetic breakpoint for *I. balthica* in the Öresund, which geographically separates the high- and low-salinity populations. This area exhibits the steepest salinity gradient of the entire Baltic Sea (including the transition zone; Johannesson et al. 2020), and can also be thought of as a separation point between the older species range and the more recently colonised area in the Baltic Sea. That the Öresund strait genetically separates the low-salinity population from the high-salinity population is further supported by a lack of population structure north of the Öresund (in the high-salinity environment), but with a strong isolation-by-distance genetic pattern observed along the Swedish coast to the southeast of the Öresund, indicative of a step-wise colonization event with multiple bottlenecks (De Wit *et al.* 2020). During a step-wise colonization history, successive locally adaptive evolution of tolerance to low salinity may have evolved during range expansion, possibly in combination with the evolution of locally adaptive plasticity (although other factors could have been involved; for example, the colonisation of *I. balthica* over new geographic areas most likely occurred after these areas were successfully colonised by *Fucus* spp.). Our simulation results (Figure S2D) also indicate some genetic differentiation in the scenario that seems most likely for the Baltic Sea population of *I. balthica*, i.e., that the species has a wide tolerance to different salinity levels (as shown by Wood et al. 2014) but that the low-salinity population has lost some of this tolerance. Some of this genetic differentiation could be due to the difference in plasticity between the two populations. However, further quantitative comparisons between the empirical and our model data are not warranted at present. Indeed, our model was built on general, yet simple grounds to provide insights into potential caveats associated with the interpretation of reaction norms in dependence on whether the trait being measured is an adaptive trait or as a fitness-indicator trait, while assessing the signatures of optimal and sub-optimal phenotypic buffering. In other words, our model was not specifically tailored to account for the detailed biology of *I. balthica*, nor for the precise heterogeneity of key environmental conditions in the Baltic Sea and the transition zone. Although salinity is the main abiotic factor that varies geographically in the Baltic Sea (Björck 1995, Johannesson et al. 2020), additional environmental variables may play a role in adaptation of *I. balthica* during its range expansion history. Among these factors are the palatability of *F. vesiculosus*, the temperature, the exposure to waves and the pH, although wave exposure and variability in pH are probably largest on the microhabitat level (Wahl et al. 2018). Adaptation to multiple environmental factors may generate signals of genetic differentiation beyond those corresponding to adaptation to the observed salinity gradient alone.

We thus conclude that the low-salinity population of *I. balthica* is probably more sensitive to increased salinity than the high-salinity population is to decreased salinity, possibly because plasticity has been lost through serial founder effects during colonisation of the Baltic Sea. Another possibility is that plasticity with respect to salinity has been reduced in the Baltic Sea (i.e., in our low-salinity population) because the salinity gradient is shallower there than in the Öresund (Johannesson et al. 2020) and/or because the low-salinity population experiences weaker fluctuations in salinity than the high-salinity population (because changes in wind and outflow from the Öresund result in larger fluctuations in salinity levels in the high-salinity environment than in the low-salinity environment; Bendtsen et al. 2009; Maar et al. 2011; see also ICES data https://www.ices.dk/data/dataset-collections/Pages/default.aspx)). A shallower local environmental gradient and/or weaker temporal fluctuations in the environmental conditions (as is the case here for the low-salinity *I. balthica* population compared to the high-salinity population) have been theoretically shown to result in lower local plasticity (Eriksson & Rafajlović 2022).

Notably, we would not have been able to conclude that the low-salinity *I. balthica* population had lower tolerance to the high-salinity environment than the high-salinity population had to the low-salinity environment if we had only measured respiration rate or only grazing activity. Moreover, we would not have been able to draw any conclusions if we had measured both traits, but observed an increase in both metabolic rate and grazing activity. In such a case, it would be unclear whether the phenotypic changes are locally adaptive or not because it would not be obvious whether the combination of increased metabolic rate and increased grazing is good or bad for the individuals unless also a more direct measure of fitness, such as fecundity or fecundity combined with survival, is included. Indeed, when individuals of *I. balthica* were exposed to elevated temperatures, Gutow *et al.* (2016) interpreted an increased metabolic rate combined with increased grazing as a locally adaptive response to warming, whereas Wood et al. (2014) interpreted the maintenance of metabolic rate in the same species as a sign of phenotypic buffering with respect to salinity.

Our conclusions would have been further strengthened if we had also measured a trait that is a more obvious fitness-indicator, such as growth, fecundity and/or offspring mortality. However, as this species grows through moulting, growth is non-linear, meaning that growth measurements taken at random time points are not directly comparable and hence difficult to interpret. Intermoult duration is a good proxy of growth (see e.g., Hemmi & Jormalainen (2004)), but the rapid decay of moults in our experiment prevented us from measuring this parameter. Future studies examining the relationship between respiration rate, feeding, intermoult duration, survival and fecundity could potentially clarify whether the observed phenotypic changes are fitness-related or not. For example, if the growth rate would have been unchanged for the high-salinity population in 8 psu, but changed for the low-salinity population in 16 psu, this, together with our results for respiration rate and grazing, would be a strong indication that the low-salinity population has reduced salinity tolerance relative to the high-salinity population.

Recall that we assumed in our simulated experiments that the populations had stabilised, which may not be the case for populations in the Baltic Sea. If the low-salinity population has recently colonised its habitat, we expect that its fitness would likely increase when transplanted to the high-salinity environment, but this is not consistent with the combination of increased metabolic rate and decreased food consumption. However, the opposite is also possible (*albeit* less likely) due to repeated founder events and potential spread of genotypes that are less-than-average adapted to their source population (surfing of maladapted alleles; Excoffier et al. 2009). Exploring this possibility further would benefit from simulations specifically tailored to include more detailed properties of the species (including the sensitivity and number of traits under selection, with or without plasticity) and the specific properties of the environment (here the Baltic Sea), including salinity, temperature, interactions with other species, and other relevant environmental parameters. We leave such an ambitious project for future work.

### Caveats and recommendations for future work

In the following, we summarise four key messages from our study.

### The interpretation of reaction norms differs depending on whether a trait is adaptive or a fitness-indicator

Naïvely interpreting reaction norms as if the assessed trait is adaptive may lead to the wrong conclusions when the assessed trait is, instead, an indicator of fitness. The same is true when the assessed trait is adaptive but wrongly assumed to be a fitness-indicator. Therefore, we suggest that researchers should consider in advance whether the traits assessed in an experiment are adaptive or fitness-indicators in the species in question and the given environmental context. However, if it is not known *a priori* whether a trait assessed is adaptive or a fitness-indicator, both possibilities should be considered along the lines illustrated in this study.

We note that it is possible that traits lie on a continuous spectrum between purely adaptive and purely fitness-indicator traits due to, for example, coupling (realised through, e.g., linkage disequilibrium or pleiotropy) between adaptive and fitness-indicator traits. This could be species and/or environmental specific.

We, therefore, encourage researchers to report whether the assessed trait is a fitness-indicator or an adaptive trait because this will help clarifying on which grounds their interpretation of the empirical results is based on. With this, it will also be possible to perform meta-analyses that will allow us to understand the type of different traits and to which extent the type of a given trait is species and/or environment specific.

### Phenotypic buffering may be sub-optimal

Recall that phenotypic buffering (*sensu* Reusch 2014) implies a flat reaction norm for a fitness-indicator trait. Here, however, we call this *optimal* phenotypic buffering, which should be distinguished from *sub-optimal* phenotypic buffering wherein reaction norms in a fitness-indicator trait are not flat. Whether phenotypic buffering is optimal or sub-optimal is likely species- and environment-dependent and is likely to vary throughout a species’ range.

Interestingly, sub-optimal phenotypic buffering can help us to distinguish between fitness-indicator and adaptive traits in reciprocal transplant experiments. Namely, if for both populations the reaction norms in a given trait have the same direction in comparison to each other, then the trait is most likely a fitness-indicator. Otherwise, if the reaction norms have the opposite direction in comparison to each other, the trait is most likely adaptive. By contrast, when optimal phenotypic buffering occurs (e.g., when the difference in optimal phenotype between native and new environment is very small) the reaction norms may be flat for both kinds of traits. In these cases, it may not be possible to tell whether the assessed trait is better described as an adaptive trait or as a fitness indicator. This may be overcome by performing additional reciprocal transplants, including locations with more distinct environmental conditions. Alternatively, if the assessed trait can reasonably be assumed to be orthogonal to other key traits of the species, one may consider the phenotypic values of the trait when optimal phenotypic buffering occurs: if the trait is adaptive, any population produces the same phenotype on average when exposed to the same environment, and if the trait is a fitness-indicator, then any population produces the same phenotype on average in all environments. Further empirical work is needed to evaluate potential candidate traits of this kind.

### Experiments conducted under laboratory conditions may not capture the effects of combined environmental variables under natural conditions

In laboratory experiments, the effect of a few environmental variables is usually studied, often without incorporating temporal fluctuations that occur in natural systems, whereas a multitude of different environmental variables typically varies in both time and space in natural habitats. This is problematic, because environmental selection under laboratory conditions can be very different from that under natural conditions. For example, survival under laboratory conditions may not imply survival in the wild and/or a surviving individual may not reproduce. In general, an adaptive response along a selection component that is studied may be maladaptive along another selection component which is absent under laboratory conditions, due to potential fitness trade-offs (e.g., Willi & van Buskirk 2022), or *vice versa*. Consequently, the inference of adaptive, maladaptive or neutral responses under laboratory conditions may not reliably inform about the responses under natural conditions. To better approximate natural populations, we encourage researchers to improve experimental designs by incorporating, in increasing complexity, multiple factors in the experiments. For example, in the case of Baltic Sea populations of *I. balthica*, experiments could be performed by accounting for variation in salinity, temperature and the geographical location from where the food (*F. vesiculosus*) is taken (due to the geographical variation in palatability of algae; Milec et al. 2022).

### Modelling and computer simulations can aid the interpretation of reaction norms

Simulations that are specifically tailored to fit the population and the environment of interest can bring further insights into adaptive responses of populations and the interpretation of reaction norms. Simulations may, for example, indicate fitness trade-offs, the extent of local plasticity and non-plastic genetic adaptation, genetic differentiation, and expectations for reaction norms with respect to the different types of traits assessed in experiments.

As a final note, we believe that more collaborative work between empiricists and theorists is necessary to gain further understanding of the role of non-plastic genetic adaptation and plasticity in the evolution of natural populations.

## Acknowledgments

We gratefully thank Kerstin Johannesson and Roger K. Butlin for insightful comments on the manuscript. We are also thankful to Per Jonsson, Luca Rugiu, and Jonathan Havenhand for valuable discussions.

This work was funded by the Hasselblad Foundation for Female Scientists (to MR), by a grant from the Swedish Research Council Formas (to MR; grant number 2019-00882), and by additional grants from the Swedish Research Councils Formas and Vetenskapsrådet, and the European Research Council through the Linnaeus Centre for Marine Evolutionary Biology (CeMEB: https://www.gu.se/en/cemeb-marine-evolutionary-biology). This work also resulted from the BONUS BAMBI project supported by BONUS (Art 185), funded jointly by the EU and the Swedish Research Council FORMAS. We would also like to thank the Swedish National Genomic Infrastructure (NGI) and the SNP&SEQ Technology platform at Uppsala University for sequencing. The computer simulations were enabled by resources provided by the Swedish National Infrastructure for Computing (SNIC) at the National Supercomputing Centre (NSC) at the University of Linköping, partially funded by the Swedish Research Council through grant agreement no. 2018–05973.

## Author contributions

MR conceived and designed the study. ME performed modelling and analyses of model data with input from MR. AK and PDW delivered and analysed empirical data. ME and PDW wrote the first draft with input from MR and AK. All authors were involved in the revisions of the manuscript.

## Conflict of interest

The authors declare no conflict of interest.

## Notes

### Competing Interest Statement

The authors have declared no competing interest.

### Summary of Updates

Added Acknowledgments and Author contributions sections. Reduced the width of Table 2.

