## Supplementary information for "Adaptive, maladaptive, neutral, or absent plasticity: Hidden caveats of reaction norms"

Martin Eriksson[1,2,3,*], Alexandra Kinnby[2,4], Pierre De Wit[2,4], Marina Rafajlović[1,2]
[1]*Department of Marine Sciences, University of Gothenburg, Gothenburg, Sweden*
[2]*Linnaeus Centre for Marine Evolutionary Biology, University of Gothenburg, Gothenburg, Sweden*
[3]*Gothenburg Global Biodiversity Centre, University of Gothenburg, Gothenburg, Sweden*
[4]*Department of Marine Sciences, University of Gothenburg, Strömstad-Tjärnö, Sweden*

### Additional details regarding the model and analyses

In this appendix, we present additional details regarding the range-expansion simulation model (see also Methods in the main text).

#### *Simulated life-cycle*

The life-cycle of the individuals during the range expansion simulations was as follows. The number of gametes that each individual contributed to the next generation was sampled from the Poisson distribution with mean equal to the individual's fitness (given by Equation (2) in the main text). Thus, the plastic response of each individual was determined by the environment where it mated. Each gamete that an individual contributed to the next generation was generated by first performing free recombination between the loci, and then choosing uniformly at random one of the homologous haploid sets of alleles to form a gamete (this process was repeated independently for each gamete that each individual contributed). For each gamete, mutation occurred symmetrically between the two possible alleles at each locus, with a uniform probability of $\mu = 10^{-6}$ per allele per gamete per generation. Among all gametes from the same deme, pairs were created uniformly at random with selfing allowed at no cost (if the total number of gametes was odd, one gamete would remain unpaired, and we discarded it). Thereafter, all adults were discarded from the simulations and the juveniles dispersed according to a discretized Gaussian dispersal function (Equations (A4)-(A6) in Eriksson & Rafajlović 2021).
Refer to the Methods section in the main text for further details regarding the range expansion model.

#### *Calculation of identity-by-state distance*

In the following, we describe how the identity-by-state distance was calculated. The genetic similarity matrix is defined as

$$\Psi = \frac{1}{L_t} ZZ^T. \tag{S1}$$

Here, $L_t$ denotes the total number of loci (for both the plastic and the non-plastic component of the trait, so that $L_t = 2L$ in the notation from Table 1 in the main text) and $T$ denotes the transpose of the matrix. The elements $Z_{k,l}$ of the matrix $Z$ (where the indices $k$ and $l$ denote individual and locus, respectively) are in turn given by

$$Z_{k,l} = \frac{G_{k,l} - p_l}{\sqrt{p_l(1-p_l)}}. \tag{S2}$$

Here $p_l$ denotes the frequency of the $+\alpha/2$ (or $+\beta/2$) allele at locus $l$ underlying the non-plastic (or plastic) component of the phenotype of the adaptive trait. The quantity $G_{k,l}$ is 0 when individual $k$ is homozygous for the $-\alpha/2$ (or $-\beta/2$) allele at locus $l$, $G_{k,l}$ is 1 when individual $k$ is homozygous for the $+\alpha/2$ (or $+\beta/2$) allele at locus $l$, and $G_{k,l}$ is 0.5 when individual $k$ is heterozygous at locus $l$.

Further details regarding the model and how simulated data were analysed are given in Methods in the main text.

### Additional results

*Linearly changing optimum*

For an optimal phenotype that changes linearly in space, that is with a constant steepness $b$ of the locally optimal phenotype (Figure S1), the gradient in plasticity was shallower than for a gradually steepening gradient in the optimal phenotype (cf. Figure 1 in the main text). However, plasticity still increased towards the edges due to, in part, increased selection for plasticity in extreme environments (Tufto 2000; Eriksson & Rafajlović, 2022).

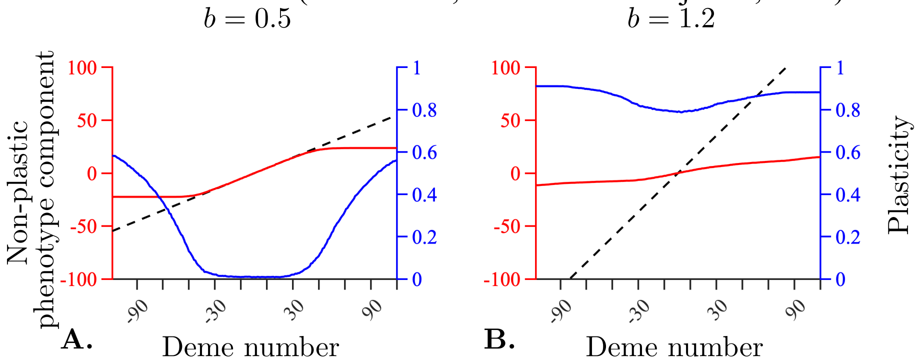


**Figure S1:** Spatial variation in the local average non-plastic component of the phenotype (red), the locally optimal adaptive phenotype (black dashed line) and local average plasticity (blue) at the end of the range expansion simulation in dependence of deme number. Shown for $b = 0.5$ (A) and $b = 1.2$ (B). To better visualise the variability in the non-plastic component of the phenotype, the $y$-axes for the locally optimal adaptive phenotype and non-plastic component are truncated at $\pm 100$. The habitat consisted of M=220 demes, numbered from -109 to 110. Remaining parameter values: $K = 100$, $r_m = 2$, $V_S = 2$, $\mu = 10^{-6}$, $L = 799$, $\alpha = \sqrt{1/10}$, $\beta = 2/L = 0.0025$, $\sigma = 1$, $\gamma = 0.5$, and $\delta = 0.5$ (Table 1 in the main text).

*Genetic differentiation between populations*

In this appendix, we present PCA plots showing differences between the two populations, accounting for all loci (underlying both the plastic and non-plastic phenotype component) in the simulated data. Figure S2 shows that the two populations from transplant experiments A, B, and C (Table 2 in the main text) have a clear genetic differentiation (Figure S2 A-C). The population from transplant experiment D is less clearly separated although all individuals from deme D2 cluster together (Figure S2 D). For transplant experiment E, the two populations do not have any obvious differentiation (Figure S2 E).

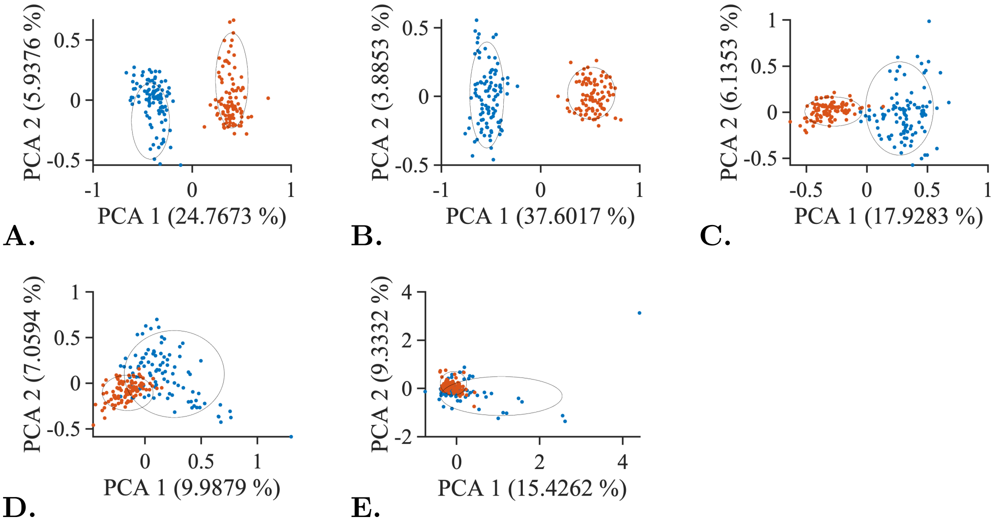

**Figure S2:** Principal component analysis of identity-by-state (IBS) distances obtained from the simulated reciprocal transplant experiments (listed in Table 2 in the main text). Individuals that are native to deme X2 are coloured in red, individuals that are native to deme X1 in blue (X=A-E in panels A-E, respectively). Ellipses represent the 95th percentiles of these two groups, respectively.

*Reciprocal transplants across longer distances*

For sampling locations spaced apart longer distances (with the two environments being A1 and B2, B1 and C2, C1 and D2, or D1 and E2), the reaction norms for the fitness-indicator trait usually had large effect sizes ($\Delta$) and were always with significant non-zero slopes (examples shown in Figure S3). In most cases, this was also true for the adaptive trait. The main exceptions were the transplants involving a population with a very low plasticity (A1 to B2), which always resulted in flat reaction norms for the adaptive trait. For transplants from B2 to A1, 996 out of 1000 realisations had non-flat reaction norms (with significant non-zero slopes), and for transplants from B1 to C2, 998 out of 1000 realisations had non-flat reaction norms for the adaptive trait. For the other transplant experiments the reaction norms for the adaptive trait were never flat. Note that for the transplant from A1 to B2, the effect sizes could still be moderately large for the adaptive trait (e.g., $\Delta = 0.21$ in Figure S3 A), even though they were not statistically significant. The two populations were typically clearly separated genetically for all reciprocal transplant experiments carried out across the longer distances (Figure S4).

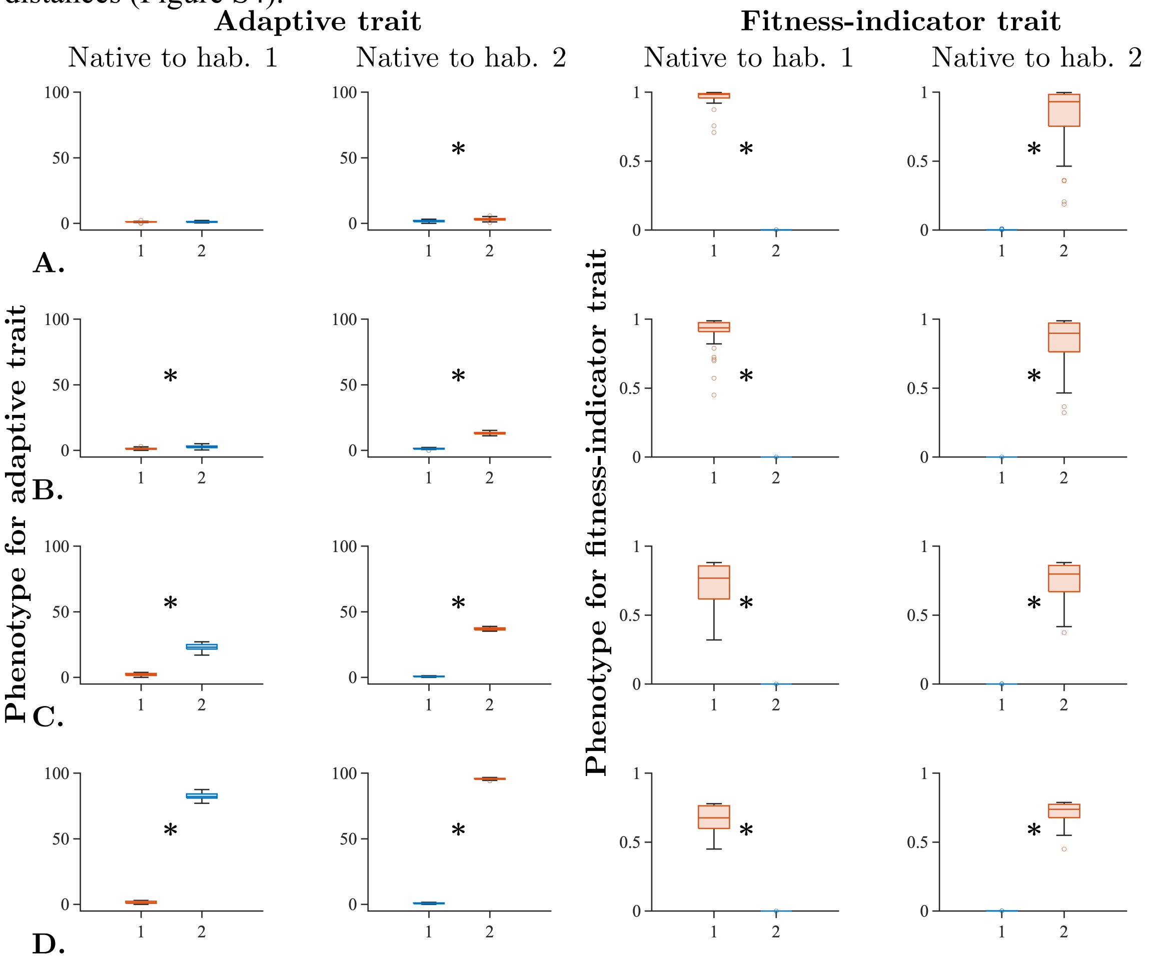


**Figure S3:** Boxplots of phenotypes from simulated reciprocal transplant experiments involving the locations denoted by A1 and B2 (row 1), B1 and C2 (row 2), C1 and D2 (row 3), and D1 and E2 (row 4) in Table 2. Measured traits are the adaptive trait $z$ (column 1 and 2) and the phenotype-dependent component of fitness (column 3 and 4). Individuals that are native to environment X1 are in column 1 and 3, individuals that are native to environment Y2 are in column 2 and 4. Here, X = A, Y=B in row 1, X=B, Y=C in row 2, X=C, Y=D in row 3, and X=D, Y=E in row 4. Kruskal-Wallis test $p$-values are: A: >0.99, $3.0 \cdot 10^{-5}$, $7.5 \cdot 10^{-30}$, $4.7 \cdot 10^{-8}$; B: $3.1 \cdot 10^{-6}$, $1.2 \cdot 10^{-24}$, $8.4 \cdot 10^{-29}$, $1.3 \cdot 10^{-8}$; C: $2.12 \cdot 10^{-5}$, $7.56 \cdot 10^{-36}$, $8.4 \cdot 10^{-26}$,$2.4 \cdot 10^{-10}$; D: $2.3 \cdot 10^{-7}$, $2.8 \cdot 10^{-31}$,

$1.0 \cdot 10^{-23}$, $9.8 \cdot 10^{-12}$. Significant ($p < 0.05$) differences are denoted with an asterisk in the plots. Glass' $\Delta$ effect sizes are: A: 0.211, -1.388, -17.569, -3.864; B: 2.384, -13.339, -7.584, -4.775; C: 22.213, -40.685, -4.950, -4.994; D: 98.715, -158.986, -6.676, -9.326. The lines inside the boxes denote the median of the data and the boxes span between the upper and lower quartiles (75th and 25th percentiles). The whiskers indicate the maximum and minimum of the non-outlier data. Outliers (denoted by rings) were computed using the interquartile range (i.e., points above the upper quartile + 1.5 times the distance between upper and lower quartile or below the lower quartile - 1.5 times the distance between the upper and lower quartile were considered outliers). Trait values in the native environments are denoted by red box plots and trait values in the new environments are denoted by blue box plots. All phenotypic values for the adaptive trait are translated so that the minimum value measured in each experiment is zero.

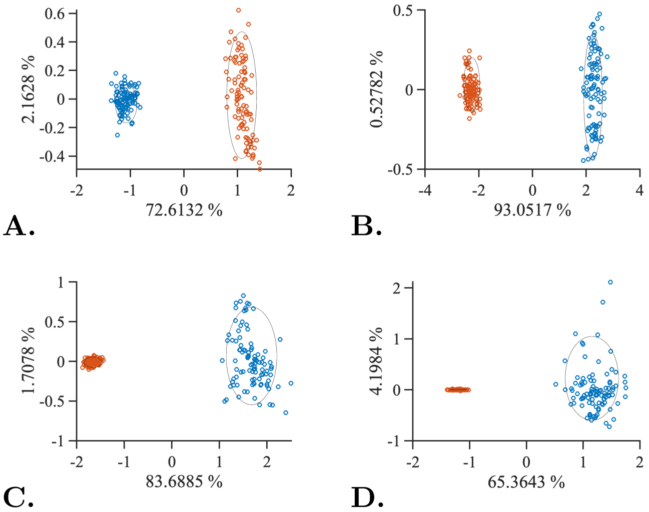


**Figure S4:** Principal component analysis of IBS distances obtained from the simulated reciprocal transplant experiments between demes A1 and B2 (A), B1 and C2 (B), C1 and D2 (C), and D1 and E2 (D). Individuals that are native to deme X2 are coloured in red, individuals that are native to deme Y1 in blue. Ellipses represent the 95th percentiles of these two groups, respectively.

*Effect sizes and significances of reaction norms as a function of local plasticity*

For a fixed difference in the phenotypic optimum $\Delta\theta$ between the two environments ($\Delta\theta = 2.5$ ), while varying the plasticity in the two environments by choosing pairs of samples at different distances from the centre (recall that plasticity is increasing towards the edges of the habitat; see Figure 1 in the main text) reciprocal transplant experiments yielded non-flat reaction norms with large effect sizes for the adaptive trait only when the local plasticity was sufficiently high (Figure S5 A, C) and for the fitness-indicator trait only when the plastic component of adaptation was sufficiently low (Figure S5 B, D).

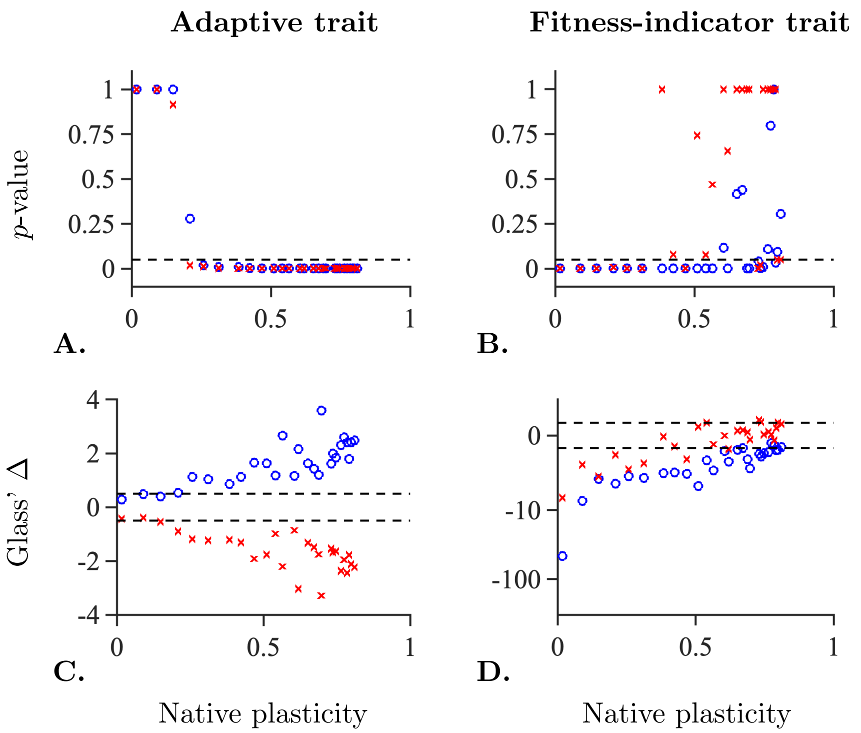


**Figure S5:** Plots of *p*-values (A-B) for non-zero slopes of reaction norms and effect sizes with direction (C-D) as a function of the locally realised average plasticity in the native environment (x-axis). The difference in optimal phenotype was kept fixed to 2.5 while the position of the two demes varied along the habitat. The blue rings stand for reaction norms for individuals that are native to the left deme (with lower plasticity) whereas red crosses stand for reaction norms for individuals that are native to the right deme (with higher plasticity). The dashed lines indicate where *p*=0.05 for the Kruskal-Wallis test (A, B), or the line beyond which absolute values of effect sizes can be considered *large* (C, D) according to Cohen's rules-of-thumb (Sawilowsky, 2009). Note a logarithmic scale on the *y*-axis in panel D.

*Effect sizes and significances of reaction norms as a function of transplant distance for a linearly changing optimal phenotype*

For optimal phenotypes that changes linearly in space, the results of reciprocal transplant experiments as a function of $\Delta\theta$ were qualitatively similar to those under a model with a steepening gradient (Figure S6; cf. Figure 3 in main text). However, because the gradient in plasticity was shallower for the linearly changing optimum, the differences between the focal and alternative environment (blue rings and red crosses, respectively) were generally smaller, making reciprocal transplant experiments with asymmetric results (i.e., significant non-zero slope for only one population) rarer compared to when the gradient was steepening. However, some asymmetry was obtained when $b = 0.5$, due to a steeper gradient in plasticity towards the edges (Figure S1 A).

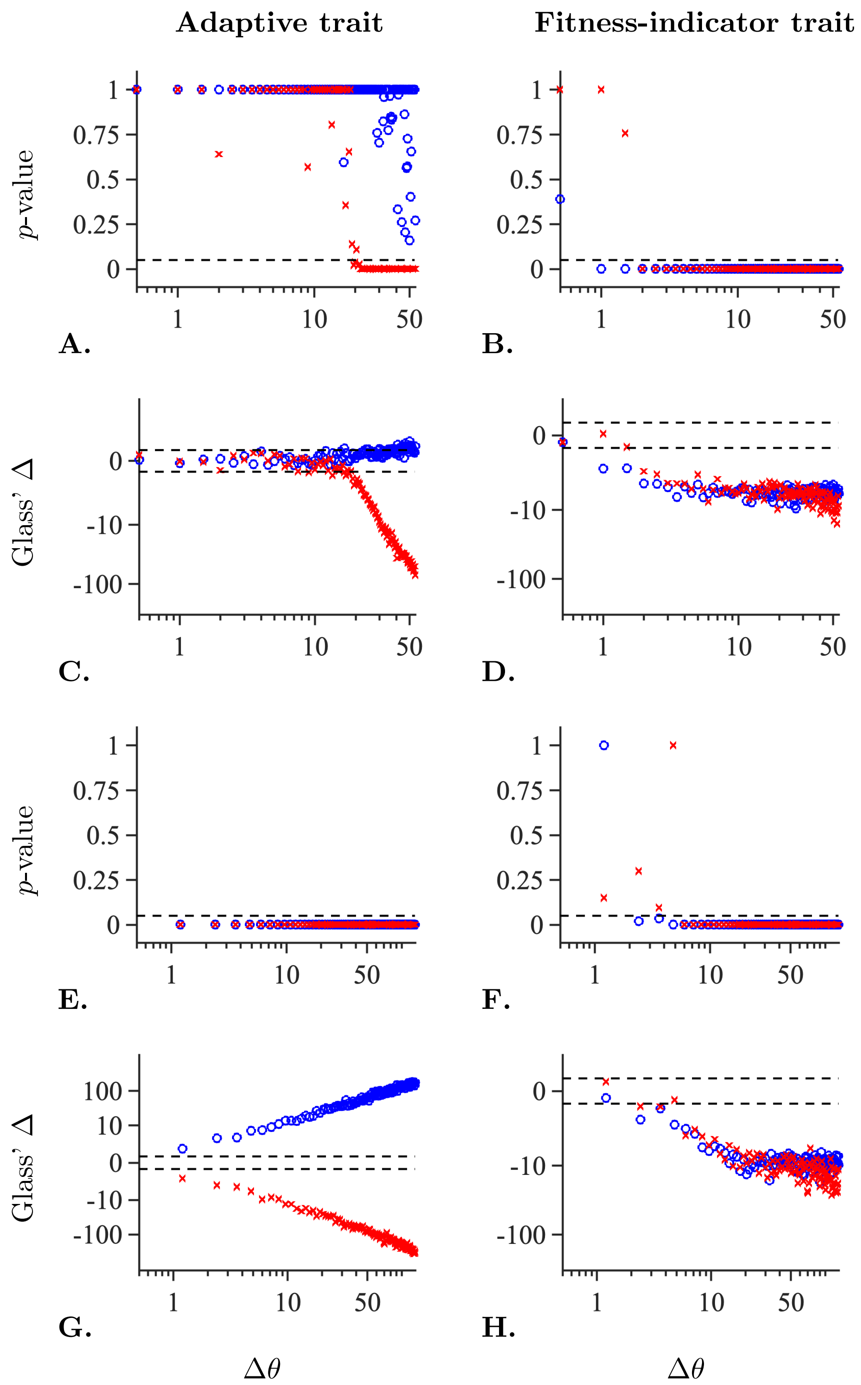


**Figure S6:** Plots of $p$-values (A, B, E, and F) for non-zero slopes of reaction norm and effect sizes with direction (C, D, G, and H) as a function of the difference in locally optimal adaptive phenotype ($\Delta\theta$) between the focal and the alternative environment for the adaptive trait (left column), or for the fitness-indicator trait (right column). Results are shown for an environmental gradient of $b = 0.5$ (A-D) and $b = 1.2$ (E-H). The dashed lines indicate where $p$=0.05 for the Kruskal-Wallis test (A, B, E, and F), or the line beyond which absolute values of effect sizes can be considered *large* (C, D, G, and H) according to Cohen's rules-of-thumb (Sawilowsky, 2009). Note a logarithmic scale on the $y$-axis in panels C, D, G, and H, and on the $x$-axis in all panels.

#### *Two-way ANOVA for empirical data*

In the following, we present the results from the two-way ANOVA analysis of the empirical data from reciprocal transplant experiments of two *Idotea balthica* (Pallas 1772) populations native to Vejbystrand (salinity of 16 psu; hereafter *the high-salinity population*) and Svarte (salinity of 8 psu; hereafter *the low-salinity population*), respectively. Two-way ANOVA was not used on the simulated data because we could not ensure normal distribution and homoscedasticity of the residuals in every realisation. The grazing rate and the respiration rate were scaled by the natural logarithm of the weight. To obtain normally distributed data, the respiration rate was log10-transformed prior to analysis. Outliers were detected using Grubb's test and removed (but results where the outliers were included are also presented).

For the grazing rate, there was no significant effect of the population, but there was a significant effect of the environment and significant gene-environment interactions (Table S1). Grazing was significantly increased in the environment with a salinity of 8 psu compared to the environment with a salinity of 16 psu, implying that *I. balthica* consumes more *Fucus* in an environment with lower salinity. Tukey's honest significance test (Tukey 1949) further reveals that a significant difference occurs between the environments with salinities of 8 and 16 psu for individuals from the low-salinity population, but not for individuals from the high-salinity population (Table S3).

For the respiration rate, there were no significant effects of the population or the environment, but a significant gene-environment interaction effect (Table S2). Tukey's honest significance test reveals that the significant differences were between the environments with salinity 8 and 16 psu for individuals from the low-salinity population, and between individuals from both populations in an environment with salinity 8 psu (Table S3).

With outliers included, there were no significant differences between the different groups for the grazing rate (Table S4). For the respiration rate, by contrast, the same qualitative results were obtained as with outliers included (Table S5).

Table S1: Two-way ANOVA for grazing rate.

**Analysis of Variance**

| Source | Sum Sq. | d.f. | Mean Sq. | F | Prob>F |
|---|---|---|---|---|---|
| X1 | 494.2 | 1 | 494.154 | 5.68 | 0.018 |
| X2 | 93 | 1 | 93.04 | 1.07 | 0.3023 |
| X1*X2 | 398.3 | 1 | 398.256 | 4.57 | 0.0335 |
| Error | 20024.5 | 230 | 87.063 | | |
| Total | 20863.7 | 233 | | | |

Constrained (Type III) sums of squares.

Note: Here, X1 is the effect of the environment, X2 is the effect of the populations. There is a significant effect of the environment, and also significant gene-environment interactions, but no significant effect of the population.

Table S2: Two-way ANOVA for respiration rate.

**Analysis of Variance**

| Source | Sum Sq. | d.f. | Mean Sq. | F | Prob>F |
|---|---|---|---|---|---|
| X1 | 0.0527 | 1 | 0.05272 | 0.73 | 0.3949 |
| X2 | 0.2747 | 1 | 0.27474 | 3.79 | 0.0529 |
| X1*X2 | 0.7848 | 1 | 0.78475 | 10.81 | 0.0012 |
| Error | 16.9077 | 233 | 0.07257 | | |
| Total | 17.9354 | 236 | | | |

Constrained (Type III) sums of squares.

Note: Here, X1 is the effect of the environment, X2 is the effect of the populations. There are significant gene-environment interactions, but no significant effects of the environment or of the population.

Table S3: Pairwise *p*-values between treatments.

| Comparison | Grazing | Respiration rate |
|---|---|---|
| **Low to low vs low to high** | 0.0156 | 0.0267 |
| **Low to low vs high to low** | 0.8560 | 0.0023 |
| **Low to low vs high to high** | 0.7635 | 0.1339 |
| **Low to high vs high to low** | 0.0841 | 0.8904 |
| **Low to high vs high to high** | 0.1226 | 0.7496 |
| **High to low vs high to high** | 0.9977 | 0.2667 |

Note: Multiple comparisons using Tukey's honest significance test reveals that the low-salinity (low) population has significant differences between the native and the new environment for both traits. The high-salinity (high) population does not have any significant differences between the native and the new environment. There is, however, a significant difference between low- and high-salinity individuals for the respiration rate in the low-salinity (8 psu) environment. Here, "low to low vs low to high" means that the low-salinity population in its native environment is compared to the low-salinity population in the high-salinity environment (and similarly for the remaining rows). These results are consistent with those obtained using the Kruskal-Wallis test followed by Bonferroni's *post hoc* test (Figure 4 in the main text).

Table S4: Two-way ANOVA for grazing rate with outliers included.

**Analysis of Variance**

| Source | Sum Sq. | d.f. | Mean Sq. | F | Prob>F |
|---|---|---|---|---|---|
| X1 | 229.8 | 1 | 229.786 | 2 | 0.1586 |
| X2 | 107.7 | 1 | 107.71 | 0.94 | 0.3339 |
| X1*X2 | 430.1 | 1 | 430.103 | 3.74 | 0.0542 |
| Error | 26656.2 | 232 | 114.898 | | |
| Total | 27320.4 | 235 | | | |

Constrained (Type III) sums of squares.

Note: Here, X1 is the effect of the environment, X2 is the effect of the populations. There are no significant effects of the environment or of the population, and no significant gene-environment interactions.

Table S5: Two-way ANOVA for respiration rate with outliers included.

**Analysis of Variance**

| Source | Sum Sq. | d.f. | Mean Sq. | F | Prob>F |
|---|---|---|---|---|---|
| X1 | 0.2249 | 1 | 0.2249 | 2.92 | 0.0888 |
| X2 | 0.1493 | 1 | 0.14933 | 1.94 | 0.1651 |
| X1*X2 | 0.4417 | 1 | 0.44169 | 5.73 | 0.0174 |
| Error | 18.0226 | 234 | 0.07702 | | |
| Total | 18.7859 | 237 | | | |

Constrained (Type III) sums of squares.

Note: Here, X1 is the effect of the environment, X2 is the effect of the populations. There are significant gene-environment interactions, but no significant effects of the environment or of the population.

These and other results we obtained are further discussed in the main text.
